# Cryptic whole-genome duplication in bryozoans

**DOI:** 10.64898/2026.09.16.752000

**Authors:** Thomas D. Lewin, Dearbhaile Casey, Yi-Jyun Luo, Anthony K. Redmond, Peter W. H. Holland

**Affiliations:** Department of Biology, University of Oxford, Oxford, UK; School of Biological and Behavioural Sciences, Queen Mary University of London, London, UK; Biodiversity Research Center, Academia Sinica, Taipei, Taiwan; School of Medicine, University College Dublin, Dublin, Ireland

**Keywords:** Paleopolyploidy, Gene duplication, Genome evolution, Invertebrates, Rediploidisation, WGD

## Abstract

Whole-genome duplications (WGDs) have long been proposed as evolutionary facilitators of biological complexity and diversification. Ancient WGDs occurred deep in the evolutionary history of both plants and vertebrates, but appear to be rare in invertebrates. However, it is unclear whether this reflects genuine scarcity or inadequate detection methodologies, and current methods may be insufficient to detect very old WGDs where extensive gene loss has followed. Here, we used a combination of paralogous and orthologous synteny-based methods incorporating bilaterian ancestral linkage groups (ALGs) to search for genomic signatures of WGD in members of the invertebrate phylum Bryozoa. This revealed strong evidence for an ancient WGD in the freshwater bryozoan *Cristatella mucedo,* Class Phylactolaemata, with duplicated paralogous regions across every chromosome. Only ∼10% of duplicated genes are retained, including a Hox cluster duplication. All eight genomes from the bryozoan class Gymnolaemata also show signatures of ancient WGD in the form of 2:1 ratios in orthologous synteny comparisons, which remain detectable despite only ∼5% duplicate retention. Gene trees support a scenario in which all extant bryozoans share an ancient WGD and much of the genome rediploidised independently in the two clades. Duplicates retained after bryozoan WGD are enriched in cilia genes and are preferentially expressed in the lophophore, the tentacular feeding organ covered in highly specialised cilia. We argue that cryptic WGD in bryozoan evolution facilitated adaptation to a sessile, filter-feeding lifestyle, and propose that incorporation of ALGs adds to the power of synteny-based WGD-detection methods.

## Introduction

Whole-genome duplication (WGD or polyploidisation) events, which create two or more complete sets of chromosomes, are considered to be major drivers of evolutionary innovation. This is because they generate duplicate copies of every gene in the genome, providing raw material from which selection can produce evolutionary novelties and increases in biological complexity^1–3^. Consistent with this, WGDs are associated with several major evolutionary radiations, for instance occurring at the base of both the seed plants and flowering plants^4–8^. Similarly, all jawed vertebrates have undergone two rounds of WGD^1,9–14^, with additional events in teleosts^15^, salmonids^16,17^, sturgeons/paddlefishes^18^, cyprinids^19–22^, and amphibians^23,24^. However, these groups appear to be the exception rather than the rule, and ancient WGDs that have persisted over long evolutionary timescales are in general extremely rare^2^.

In stark contrast to plants and vertebrates, very few ancient WGDs have been described in invertebrates and WGDs of any age are currently known from only six of ∼32 animal phyla^25^ (Fig. 1A). Invertebrates with described WGDs are restricted to clitellates^26–28^ (phylum Annelida); chelicerates^29–33^ and barnacles^34^ (Arthropoda); caenogastropods^35,36^, stylommatophorans^37,38^, and bivalves^39^ (Mollusca); the rhabditid *Allodiplogaster sudhausi*^40^ and three species of *Meloidogyne* root-knot nematodes^41^ (Nematoda); *Macrostomum* flatworms^42,43^ (Platyhelminthes), and rotifers^44–46^ (Rotifera). Many of these WGDs appear to be relatively recent and are restricted to small clades, but it is not clear whether the apparent scarcity of ancient invertebrate WGDs is genuine or an artefact caused by challenges in detection, especially in taxa with fast-evolving and divergent genomes. In particular, WGD detection methods such as synonymous substitution (*K_s_*) distributions^47–49^ and gene tree-species tree reconciliation^7,50–52^ are prone to failure in cases of extensive gene loss following WGD. In contrast, recent work using paralogous (within-genome)^53^ and orthologous (cross-species) synteny has been successful in disentangling ancient WGDs in the flowering plant^8,53,54^ and vertebrate^9,10,55^ lineages. Cross-species analyses are especially valuable for ancient WGDs because WGD signatures can be detected even if most duplicate genes (ohnologues) have returned to single-copy, as this would still leave a signature of ‘double conserved synteny’: 2:1 ratios where dispersed genes on two chromosomes match their orthologues on one chromosome in an outgroup^56^. We also reasoned that the strong conservation of bilaterian ancestral linkage groups (ALGs) in many invertebrate clades^57,58^—gene sets kept together on the same chromosome since the ancestor of a given species set^13,57^—would give us additional power to detect this^59–61^.

**Fig. 1.**
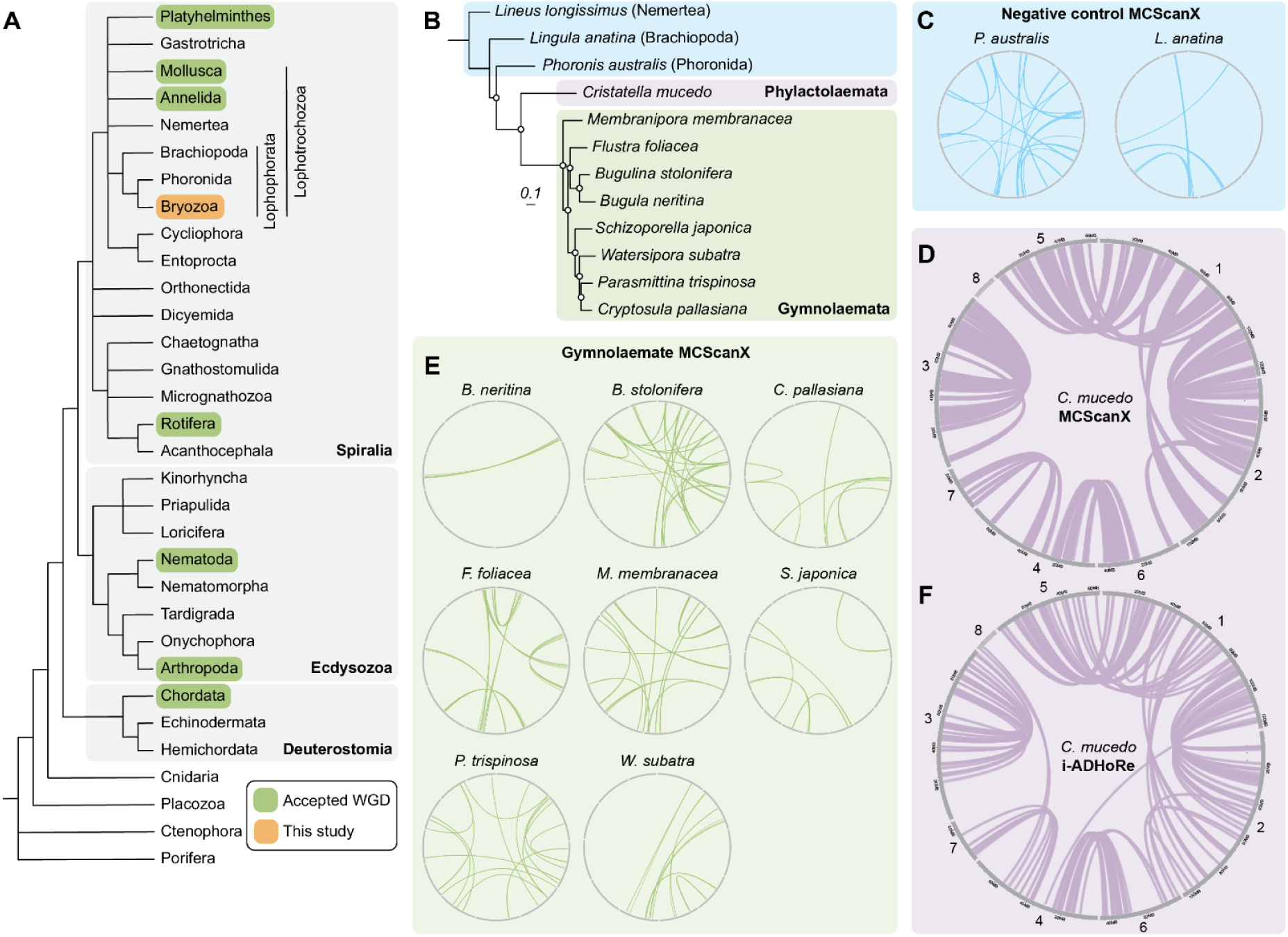
Intra-genomic synteny suggests a WGD in the bryozoan *Cristatella mucedo*. A. Tree of animal phyla adapted from^109^ showing lineages with known WGD events. B. Tree of bryozoan genomes used in this study. The tree was constructed using the maximum likelihood method of IQ-TREE^110^ using 1879 single-copy orthologues (963,051 amino acid positions). The Q.PFAM+F+R7 model was selected by ModelFinder^111^ as the optimum sequence evolution model. Branch lengths represent substitutions per site. White circles at nodes indicate maximum support. C. Circos plot showing the distribution of duplicated collinear gene blocks (paralogons) in two negative control genomes, as determined by MCScanX^107^. Grey bars are chromosomes. Blue lines connect the position of paralogons. Only cross-chromosome paralogons are shown. D. Circos plot showing the distribution of paralogons in the *C. mucedo* genome, as determined by MCScanX^107^. Grey bars are chromosomes. Purple lines connect the position of paralogons. Only cross-chromosome paralogons are shown, with the exception of chromosome 3. E. Circos plot showing the distribution of paralogons in the eight gymnolaemate bryozoan genomes, as determined by MCScanX^107^. Grey bars are chromosomes. Green lines connect the position of paralogons. Only cross-chromosome paralogons are shown. F. Circos plot showing the distribution of paralogons in the *C. mucedo* genome, as determined by i-ADHoRe^108^. Grey bars are chromosomes. Purple lines connect the position of paralogons. Only cross-chromosome paralogons are shown, with the exception of chromosome 3.

One group of animals in which the presence of WGD has not been detected is the phylum Bryozoa, also known as ectoprocts or moss animals. Most bryozoans are colonial, sessile, filter-feeders found in marine or freshwater habitats (Fig. 1B). Individual zooids each have their own mouth, gut, nervous system, and lophophore—a ciliated tentacle used for filter-feeding—but form colonies of typically hundreds of interconnected individuals^62–64^. Around 6,500 extant species have been described to date^65^, supplemented by a rich fossil history that stretches back to the Cambrian^66–70^. While globally-distributed and ecologically important, they are understudied compared to other lophotrochozoans such as annelids and molluscs^69,71^. However, recent years have seen the increasing application of molecular techniques to bryozoans, including molecular phylogenetics^72–84^, transcriptomic profiling^85–88^, and recently the production of high quality chromosome-level genome assemblies^89–96^.

In previous genomic work, we briefly postulated a bryozoan WGD but considered it unlikely as *K_s_*distributions showed no temporal peak in rate of gene duplications^95^. However, this, and other widely-used approaches, may miss hard to detect WGDs, such as those that are particularly ancient or those followed by extensive loss of gene duplicates^49,97,98^. In this work, we used newly-released chromosome-level genomes and a comparative genome structure-based approach incorporating paralogous synteny, orthologous synteny, and bilaterian ALGs, to test whether some or all bryozoans underwent an ancient WGD event.

## Results

### Intra-genomic synteny reveals WGD in the bryozoan *Cristatella mucedo*

To test for possible ancient genome duplications, one can use either comparisons within a single genome (paralogous synteny) or comparisons between species (orthologous synteny). Within a single genome, strong evidence for WGD would be provided by the presence of many duplicated collinear gene blocks (paralogons) distributed throughout^99–102^. Immediately post-WGD, these would be expected to span the length of each pair of duplicated chromosomes. Over time, rearrangements and gene losses, which can occur quickly due to functional redundancy between duplicated genes, shuffle and erode these regions, reducing them to smaller blocks and making them more difficult to detect. To test whether a paralogon signature is present in bryozoans, we first obtained all available chromosome-level genomes (*n* = 9) and annotated those without published gene models using BRAKER3^103^. Extant bryozoans are split into three classes: Phylactolaemata, Stenolaemata, and Gymnolaemata. Our dataset consisted of one phylactolaemate (*Cristatella mucedo*) and eight gymnolaemates, covering the deepest split in living bryozoans (Fig. 1B, Supplementary Table 1). We also added representatives of bryozoans’ two closest relatives, phoronids and brachiopods, with which they form the clade Lophophorata^75,104–106^.

We searched for paralogous synteny in Lophophorata genomes using the collinearity detection algorithm MCScanX^107^. Very few paralogons are identified in the negative controls, *Phoronis australis* (Phoronida) and *Lingula anatina* (Brachiopoda) (Fig. 1C). By contrast, the phylactolaemate bryozoan *C. mucedo* has paired paralogons distributed across its genome (Fig. 1D). Large stretches of collinear blocks are present between chromosomes 1 and 2, chromosomes 1 and 5, chromosomes 4 and 6, chromosomes 4 and 7, and the two halves of chromosome 3, which is highly suggestive of WGD. Only chromosome 8 does not show clear homology to another chromosome. This suggests that a WGD occurred in the evolutionary history of *C. mucedo* followed by limited subsequent rearrangement, preserving blocks of ohnologues. Unlike *C. mucedo*, we detect only very few paired paralogons across the eight gymnolaemate genomes (Fig. 1E) and duplicate gene distributions are similar to the negative controls *P. australis* and *L. anatina* (Fig. 1C). We return to this finding later in this manuscript.

We next checked these results with an independent collinearity search algorithm, i-ADHoRe^108^. This corroborated the presence of paralogons throughout the genome of *C. mucedo* (Fig. 1F) but not gymnolaemate bryozoans, phoronids, or brachiopods (Supplementary Fig. 1). In *C. mucedo*, i-ADHoRe supports the pairs of chromosomal homologies suggested by MCScanX and additionally finds paralogons shared between chromosome 8 and chromosome 4 (Fig. 1F). Overall, paralogous synteny strongly supports a WGD in the phylactolaemate *C. mucedo*.

### Cross-species synteny is compatible with WGD in gymnolaemate bryozoans

The proportion of duplicated Benchmarking Universal Single-Copy Orthologs (BUSCO)^112^ genes in the *C. mucedo* genome is only 4%, suggesting extensive gene loss post-WGD and the return of most genes to single copy (Supplementary Fig. 2, Supplementary Table 2). BUSCO duplication counts in gymnolaemates are even lower, ranging from 1.6 to 3.4%. Although BUSCO genes cannot be used to estimate genome-wide gene retention, we considered the possibility that gymnolaemates too have undergone a WGD but higher rates of gene loss obscured any paralogon signal beyond the limits of detection. To test this, we needed a WGD detection method that does not rely on blocks of duplicate genes. In doubling every region in the genome, a WGD creates a 2:1 mapping of genomic regions compared to close outgroups without a WGD (Supplementary Fig. 3). In theory, this 2:1 ratio should remain detectable even if all duplicates return to single copy, provided no extensive interchromosomal rearrangements have taken place. Cross-species synteny-based analyses (or orthologous synteny^8,54^) that reveal such correspondences could therefore offer a clear test for WGD^8,56,113–115^. We also considered that the incorporation of data from ancestral linkage groups (ALGs)^13,57^ might increase robustness to chromosomal rearrangements. Each bilaterian ALG is a set of genes co-located on the same chromosome in the ancestor of bilaterian animals^57^, and is taken as a representative of ancestral chromosomes. In the absence of WGD, we expect the majority of genes from each ALG to be found on a single chromosome.

We began by using dot plots of single-copy orthologous genes from the 24 bilaterian ALGs to compare the structure of the nine bryozoan genomes to that of the phoronid *P. australis*, their nearest outgroup^75,105,106,116,117^ (Fig. 2A). *P. australis* has undergone only minimal rearrangements to the bilaterian ALGs and each of the 24 is found only on a single chromosome^75^, confirming that there has been no WGD in its history and making it a suitable model for comparison with bryozoans. In gymnolaemate bryozoans, 23 of the 24 ALGs are found on either two or four chromosomes. For instance, in *M. membranacea*, ALGs A2, B1, B2, B3, C1, D, E, G, H, I, J1, J2, L, O1, Q, and R are found on four chromosomes (chromosomes 1, 2, 3, and 4) (Fig. 2A). ALGs A1 and C2 (chromosomes 6 and 8), F and N (chromosomes 5 and 11), and K, O2, and P (chromosomes 9 and 10) are found on two chromosomes. Each of these ALGs maps to only one chromosome in *P. australis*, leaving 2:1 and 4:1 ratios (Fig. 2A). ALG M is an outlier, being dispersed across many chromosomes. These 2:1 and 4:1 ratios, representative of ancestral chromosomes, are suggestive of either WGD or many chromosome fissions in *M. membranacea*.

**Fig. 2.**
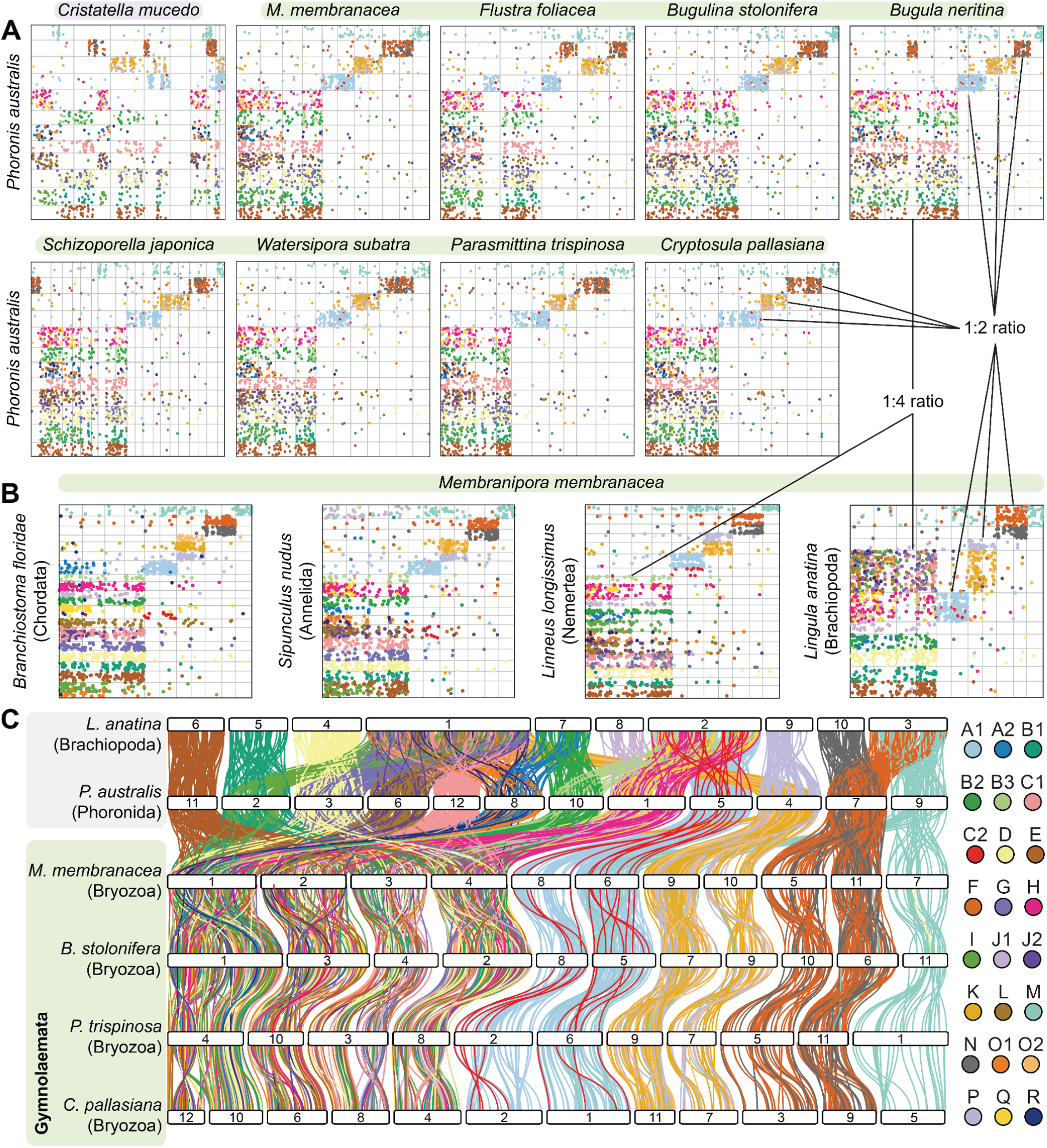
Cross-species synteny supports WGD or multiple fissions in gymnolaemates. A. Dot plots of bryozoan genomes with that of their closest relative, phoronids, represented by *Phoronis australis*. Each plot represents a pairwise comparison of two genomes: one bryozoan (X-axis) and *P. australis* (Y-axis). Each axis represents the full length of the genome with chromosomes laid end-to-end. Grey bars mark chromosome boundaries. Each dot marks the position of a single-copy orthologous gene in the two genomes. Dots are organised into quadrangles, representing groups of genes that have been maintained together on the same chromosome in both genomes i.e. the conservation of macrosynteny. Dots are coloured by their bilaterian ancestral linkage group—dots of the same colour were on the same chromosome in the ancestor of bilaterians. Across all gymnolaemate bryozoans (all except *C. mucedo*), a 1:2 or 1:4 ratio is observed between phoronid and bryozoan chromosomes. This is suggestive of one or two rounds of WGD. B. Dot plots of the bryozoan *M.* with representatives of the phyla Brachiopoda, Annelida, Mollusca and Chordata. In each case, a 1:2 and 1:4 ratio of chromosomes is observed. C. Riparian plot for selected gymnolaemates and the nearest outgroup to bryozoans, phoronids and brachiopods. White bars represent chromosomes. Vertical ribbons connect the position of orthologous genes, coloured by their bilaterian ALG. In outgroup phyla, each ALG is found on one chromosome. In most gymnolaemate bryozoans, each ALG is found on two (A1, C2, F, K, N, O2, P) or four (A2, B1, B2, B3, C1, D, E, G, H, I, J1, J2, L, O1, Q, R) chromosomes. This is suggestive of either one or two rounds of WGD or one or two fissions of each ancestral chromosome. Riparian plot including all species is available as Supplementary Fig. 6.

Four other gymnolaemates *B. neritina*, *B. stolonifera*, *P. trispinosa*, and *W. subatra* share this same set of 2:1 and 4:1 ratios when compared with *P. australis*. A further two show only slight deviations: *F. foliacea* (2:1 and 3:1 ratios) and *C. pallasiana* (2:1 and 5:1 ratios). *S. japonica* is the one outlier, with 3:1 (ALGs K, O2, and P), 4:1 (ALGs A1, C2, F and N), and 8:1 (ALGs A2, B1, B2, B3, C1, D, E, G, H, I, J1, J2, L, O1, Q, and R) ratios. Further interrogation shows that this species specifically has undergone lineage-specific fissions and fusions resulting in an unusual karyotype that complicates analysis but is also compatible with a single WGD (Supplementary Discussion 1, Supplementary Fig. 4, 5, Supplementary Table 3). The phylactolaemate bryozoan *C. mucedo* has undergone more substantial rearrangement to the bilaterian ALGs^95^, preventing clean cross-species comparisons (Fig. 2A). In summary, gymnolaemate bryozoan genomes in general have a 2:1 and 4:1 ratio of bilaterian ALGs (representing ancestral chromosomes) compared to those of their nearest outgroup, the phoronids.

We then questioned whether these 2:1 and 4:1 ratios were a quirk of the comparison with phoronids—for instance, due to lineage-specific chromosome fusions in phoronids. We compared the genome of *M. membranacea* (representing gymnolaemates) to representatives from four further phyla with relatively conserved genomic structures^58^: *Lingula anatina* (Brachiopoda), *Lineus longissimus* (Nemertea), *Sipunculus nudus* (Annelida), and the more distantly related *Branchiostoma floridae* (Chordata) (Fig. 2B). In each case, the 2:1 and 4:1 chromosomal ratios were preserved, confirming that these ratios are due to either the duplication or fission of chromosomes in gymnolaemate bryozoans.

Dot plots are useful for observing ratio changes but obscure specific details and make comparisons across more than two species difficult. We therefore used single-copy orthologs as markers of chromosomal orthology and visualised their positions across the genomes of bryozoans and representative outgroups using riparian plots (Fig. 2C, Supplementary Fig. 6, 7). This revealed strong conservation of genome structures across the gymnolaemates, with the 11 chromosomes described previously^95^ conserved in four out of eight species. It also illuminated the cause of the deviations from a 4:1 ratio in several species: the 5:1 ratio between *C. pallasiana* and outgroups is caused by the fission of one chromosome (now chromosomes 10 and 12), leaving 12 chromosomes in total (Fig. 2C), and the 3:1 ratio between *F. foliacea* and outgroups is due to a fusion event of two chromosomes containing the same ALGs to form chromosome 1, consistent with a previous report^96^. Fusions of chromosomes carrying different ALG sets do not affect ratios: for instance, *B. neritina* has two fusions of chromosomes carrying different ALGs, but maintains the 2:1 and 4:1 ratios. Overall, the observed 3:1 and 5:1 ratios are 4:1 ratios modified by derived lineage-specific fusions or fissions. Therefore, orthologous synteny incorporating ALGs reveals that gymnolaemate bryozoans have a clear 2:1 and 4:1 ratio of chromosomes compared to other related phyla. This points towards one of three scenarios: one gymnolaemate WGD, two gymnolaemate WGDs, or many chromosome fissions.

### Enrichment of gene duplicates supports a gymnolaemate WGD

The observed 2:1 and 4:1 chromosome homology ratios may be best explained by WGD but we must first rule out the possibility that they are the product of chromosome fissions. These two hypotheses make different predictions for the distribution of gene duplicates. The expectation in the case of WGD is that duplicated genes will be present on paralogous chromosomes within a given genome^114^. This produces pairs of chromosomes with an enrichment of shared duplicates (Fig. 3A), even if intra-chromosomal rearrangements have eliminated conserved gene order such that paralogons become difficult or impossible to detect. In contrast, chromosome fission is not expected to produce pairs of chromosomes with an enrichment of duplicate genes, especially on a genome-wide scale (Fig. 3B).

**Fig. 3.**
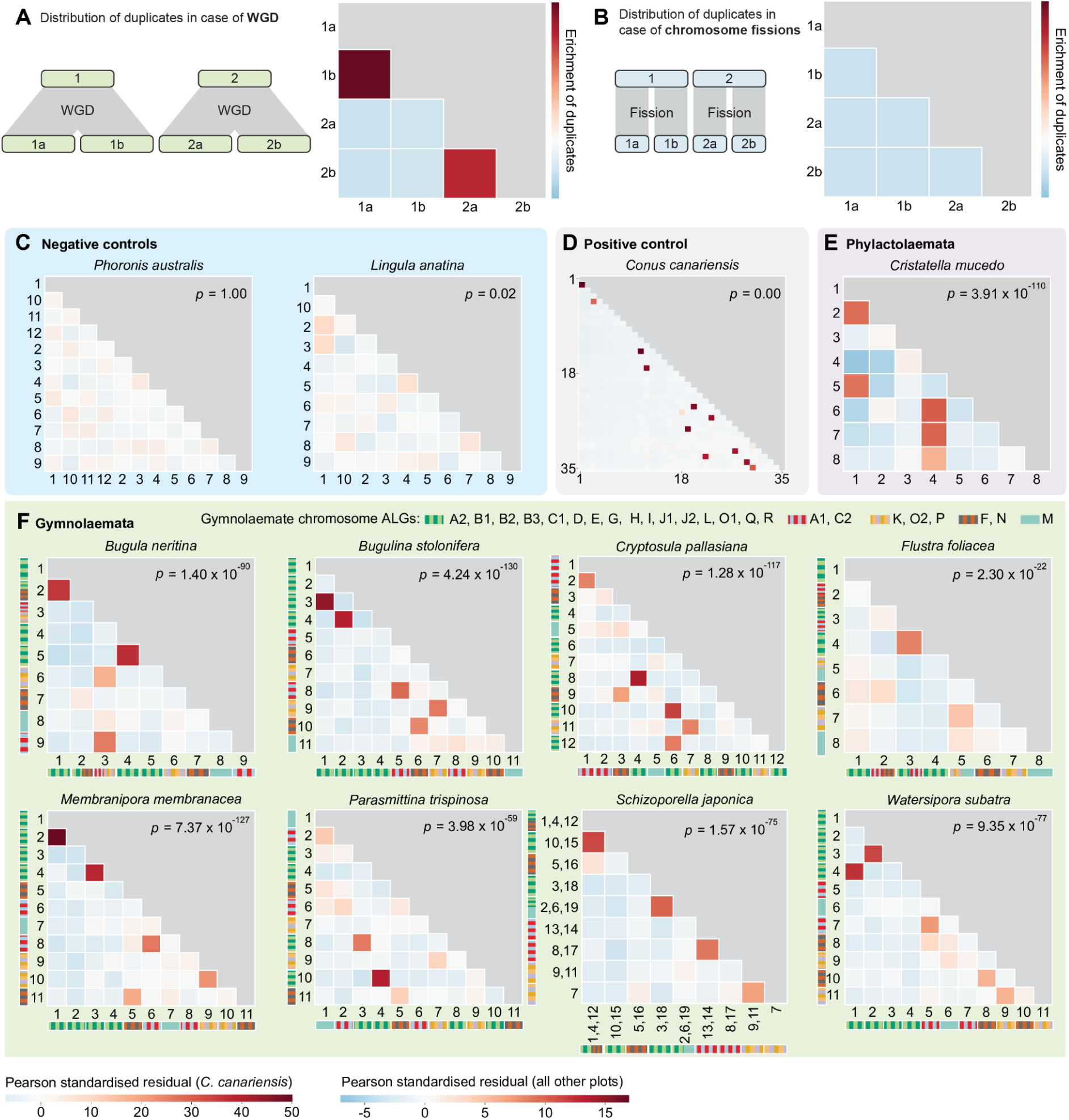
Enrichment of duplicated genes on putative ohnologous chromosomes supports WGD over fissions. A. Expectations for enrichment of duplicate pairs when chromosomes are the product of WGD. B. Expectations for enrichment of duplicate pairs when chromosomes are the product of fission events. C–F. Heatmaps showing chromosome pairs enriched (red) and depleted (blue) in duplicate gene pairs. Each chromosome pair is represented by one square. Coloured bars show ALG composition of Gymnolaemate chromosomes. Putative ohnologous chromosomes in bryozoans are the most enriched in duplicated genes (warmer colours). For instance, in *M. membranacea*, the most enriched pairs are chr1/chr2, chr3/chr4, chr6/chr8, chr9/chr10, and chr5/chr11. Global Bonferroni-corrected chi-squared *p*-values are shown. All bryozoans and the positive control have highly significant *p*-values while the negative controls are non-significant or weakly significant. *S. japonica* chromosome pairs derived from fission were fused *in silico* to demonstrate that this species shows the same pattern as other gymnolaemates despite its divergent karyotype.

To test which pattern is found in bryozoan genomes, we identified duplicate gene pairs and mapped their chromosomal locations, then tested for a non-random distribution of duplicates between chromosome pairs using chi-square tests. Where the global test indicated a significant departure from random pairing, standardised Pearson residuals were used to identify chromosome pairs contributing most strongly to this signal. We first validated that this method could distinguish fissions from duplication using known events (Supplementary Discussion 2, Supplementary Fig. 8, 9) and then used positive and negative controls for further validation (Fig. 3C). Negative controls (phoronids and brachiopods with no WGD) showed no strong enrichment of duplicates across any pair of chromosomes (*P. australis*, *p* = 1.00; *L. anatina*, *p* = 0.02) (Fig. 3C). *Conus canariensis*, a positive control with a recent WGD, showed a strong enrichment of duplicates (*p* = 0.00) and clear pairs of ohnologous chromosomes (Fig. 3D).

In *C. mucedo*, where our paralogon analyses identified a clear WGD, duplicates are very strongly enriched (*p* = 3.91 x 10^-110^) across the chromosome pairs identified with MCScanX and i-ADHoRe: chromosomes 1 and 2, chromosomes 1 and 5, chromosomes 4 and 6, chromosomes 4 and 7, chromosomes 4 and 8 (Fig. 3E). This confirms the proposed WGD and validates the method.

All gymnolaemate bryozoans also showed non-random dispersal of duplicates and strong enrichments across several chromosome pairs (Fig. 3F, Supplementary Table 4). For instance, in *M. membranacea*, strong enrichments of duplicates are found between chromosomes 1 and 2, chromosomes 3 and 4, chromosomes 5 and 11, chromosomes 6 and 8, and chromosomes 9 and 10 (Fig. 3F). Crucially, these are the same chromosome pairs identified as putative WGD-derived homologues by the cross-species synteny analysis. This implies that these chromosomes are the product of WGD and not fission, because they possess an enrichment of duplicate genes. Moreover, this is the case for all eight of the gymnolaemate genomes (Fig. 3F), suggesting that all gymnolaemates analysed have a WGD.

This analytical framework also allows us to distinguish between one and two WGD events in gymnolaemates. Although some ALGs show a 4:1 ratio in gymnolaemates when compared to outgroups (Fig. 2A, B, e.g. *M. membranacea* chromosomes 1, 2, 3, and 4), enrichment of gene duplicates reveals this actually reflects two pairs, rather than a quartet. Duplicates are enriched between *M. membranacea* chromosomes 1 and 2, and chromosomes 3 and 4, but not between other combinations (1 and 3, 1 and 4, 2 and 3, 2 and 4) (Supplementary Fig. 8). This pattern is found in all gymnolaemates (Fig. 3F), and is consistent with a single WGD in the evolutionary history of gymnolaemate bryozoans. Overall, a combination of orthologous synteny and pairwise chromosomal enrichment of gene duplicates demonstrates that extant gymnolaemate chromosomes are the product of one WGD event and not chromosome fissions.

### Both phylactolaemate and gymnolaemate bryozoans have a duplicated Hox cluster

Hox clusters are frequently retained in duplicate post-WGD but rarely duplicated without WGD, meaning the possession of multiple clusters is a signature of WGD^29,118–120^. Previous work has shown that both gymnolaemates and phylactolaemates have Hox genes on two chromosomes^121^ (Supplementary Fig. 10), but how these relate to Hox cluster duplication has not been tested. In particular, it could be argued that presence of only a single Hox gene on one of the gymnolaemate chromosomes may be a remnant of a duplicated cluster but could instead be the product of a single gene duplication. We searched for duplicated flanking genes next to the putative Hox clusters to distinguish between full cluster duplications (consistent with WGD) and single gene duplications (not the product of WGD). Duplicated flanking genes were found within 100 kb of both Hox regions in both the phylactolaemate *C. mucedo* (*nedd4* and *unc5020*) and gymnolaemates such as *M. membranacea* (*bscl2*) (Fig. 4A). This confirms that these are genuine remnants of Hox cluster duplications, as would be expected if derived from WGD events.

**Fig. 4.**
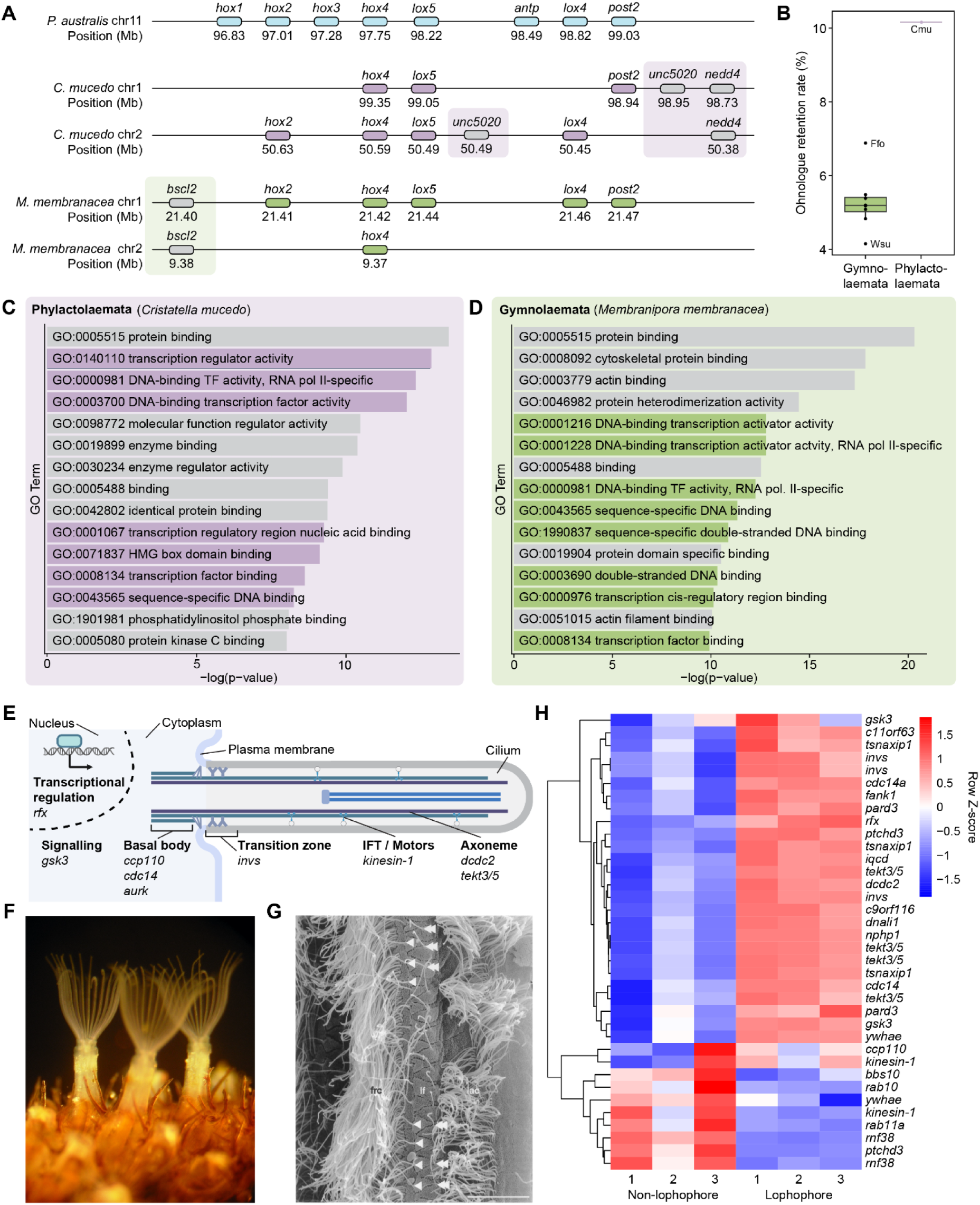
WGD-derived duplicates are enriched in transcription factors and ciliary genes. **A.** Flanking genes confirm the duplication of bryozoan Hox clusters. In the phylactolaemate *C. mucedo*, duplicated non-Hox genes *unc5020* and *nedd4* flank the two Hox clusters, acting as markers of a full Hox cluster duplication. In the gymnolaemate *M. membranacea*, the duplicated non-Hox gene *bscl2* flanks the two Hox clusters, acting as a marker of a full Hox cluster duplication. B. Estimated rate of ohnologue retention in phylactolaemate and gymnolaemate bryozoans. C. Top 15 molecular function GO terms for retained ohnologues in the phylactolaemate *C. mucedo.* Terms related to transcription factor functions are highlighted in purple. D. Top 15 molecular function GO terms for retained ohnologues in the gymnolaemate *M. membranacea.* Terms related to transcription factor functions are highlighted in green. E. Diagram of a cilium with categories of genes duplicated in both phylactolaemate and gymnolaemate bryozoans. Created in BioRender. F. Lophophores and spines of the gymnolaemate bryozoan *Flustrellidra hispida*. By Ar rouz - CC BY-SA 4.0, https://commons.wikimedia.org/w/index.php?curid=42673506. G. Scanning electron microscope image of the laterofrontal side of a lophophore tentacle of *F. hispida*, reproduced with permissions from Nielsen (2005)^138^. frc, Frontal cilia; lac, lateral cilia; lf, laterofrontal cells; single arrowheads, bases of laterofrontal cilia of the frontal row; double arrowheads, bases of laterofrontal cilia of the lateral row; scale bar, 10 µm. H. Heatmap showing expression levels of ciliary genes with multiple WGD-derived copies in lophophore versus non-lophophore tissue. Expression data is from the gymnolaemate bryozoan *F. hispida*. Colours represent row Z-score (scaled log₂ (TMM + 1) expression).

### Massive post-WGD gene loss and selective retention of transcription factors and cilia genes

We next measured the proportion of WGD-derived duplicates that have been retained as ohnologues in extant bryozoan genomes using genes that are single-copy in phoronids and brachiopods as a baseline. In *C*. *mucedo*, approximately 10.2% of genes have been retained in duplicate (Fig. 4B, Supplementary Table 5). In gymnolaemates the retention rate is even lower, on average 5.3% (Fig. 4B, Supplementary Table 5). This may explain our ability to detect collinear gene blocks with MCScanX and i-ADHoRe in the genome of *C. mucedo* but not those of gymnolaemates, which have retained half as many genes in duplicate. Overall, we approximate that 90–95% of genes have returned to single-copy following WGD in bryozoans.

Extensive gene loss places extra emphasis on the few genes that persist in duplicate. Why have these specific genes been maintained as multiple copies while the vast majority of duplicates were dispensed with? Gene ontology (GO) analysis reveals that, in the phylactolaemate *C. mucedo*, retained duplicates are strongly enriched for transcription factors (Supplementary Tables 6–9). Seven of the top 15 enriched GO terms relate to transcription factor activity and DNA binding (Fig. 4C). In gymnolaemates like *M. membranacea*, eight of the top 15 GO terms enriched in retained pairs of putative ohnologues also relate to transcription factor activity (Fig. 4D, Supplementary Tables 10–13). Transcription factor-encoding genes retained as WGD duplicates in both bryozoan lineages include *elf*, *foxn1/4*, *jun* (and, in gymnolaemates, its dimerisation partner *fos*, with which it forms the AP-1 complex^122,123^), *ror*, *rfx*, *soxc*, *tcf*, and *yy1*.

We noticed that many genes related to cilia development and function are also present in the lists of retained ohnologues. We therefore tested whether ciliary genes, as defined by the ciliary gene database CiliaCarta^124^, are enriched in the retained ohnologue sets. Strong enrichments were found in both the phylactolaemate *C. mucedo* (Fisher’s exact test *p* = 4.18 x 10^-3^, odds ratio = 1.59) and gymnolaemates, represented by *M. membranacea* (Fisher’s exact test *p* = 3.30 x 10^-5^, odds ratio = 2.42) (Supplementary Tables 14–15). Ciliary genes retained in duplicate in both clades of bryozoans include *rfx*, an ancient transcriptional regulator of ciliogenesis^125–128^; *ccp110* (the gene for CP110), an important structural regulator of ciliary assembly^129,130^; the transition zone protein *invs*^131^; *aurk*, which induces ciliary disassembly^132,133^; and *tektin3/5,* a coiled-coil domain-containing protein located in the ciliary axoneme in everything from algae to animals^134,135^ (Fig. 4E). *Tektin3/5* is of particular interest because spiralians typically have two copies^136^ and both are retained in duplicate following WGD in gymnolaemate bryozoans.

We hypothesised that ciliary genes were preferentially retained because of the critical importance of cilia on the bryozoan lophophore to generating feeding currents (Fig. 4F, G)^137^. To test this, we used an RNA-sequencing dataset^75^ to compare gene expression in lophophore and non-lophophore tissue in the gymnolaemate bryozoan *Flustrellidra hispida* (Supplementary Tables 16–18). Just over half of putative ciliary ohnologues in *F. hispida* were found to be differentially expressed between the two tissues. Importantly, 93% of these were preferentially expressed in the lophophore and only 7% show higher expression in non-lophophore tissue (Fig. 4H). All four *tektin3/5* ohnologues are upregulated in the lophophore while other genes, such as *ywhae*, appear to have undergone subfunctionalisation, with one ohnologue preferentially expressed in the lophophore and one in non-lophophore tissue. Overall, we contend that ciliary genes were retained at an elevated rate post-WGD because they were deployed in bryozoans’ ciliated feeding organ, the lophophore.

### A single WGD shared by all bryozoans?

There are two possible explanations for the presence of a WGD in members of both Phylactolaemata and Gymnolaemata. First, and most parsimoniously, a single WGD occurred in the ancestor of all bryozoans and is shared by the two clades (Fig. 5A). Second, WGD occurred in each lineage independently (Fig. 5B). To distinguish between these scenarios, we first looked at whether similar genes have been retained in duplicate in the two lineages. There is a very high degree of similarity in the duplicate gene sets retained in different gymnolaemates (e.g. 87% overlap between *C. pallasiana* and *M. membranacea*), confirming a WGD shared by all of the gymnolaemates sampled in this work. In contrast, there is much less overlap between ohnologue pairs retained in gymnolaemates and *C. mucedo*: only 31% of *M. membranacea* ohnologues are also retained in duplicate in *C. mucedo*, and only 18% of *C. mucedo* ohnologues are also retained in duplicate in *M. membranacea* (Fig. 5C). Gymnolaemates and phylactolaemates diverged over 500 million years ago (MYA)^79^ and this may therefore reflect either independent WGDs or a single, shared WGD with independent losses occurring in each lineage.

**Fig. 5.**
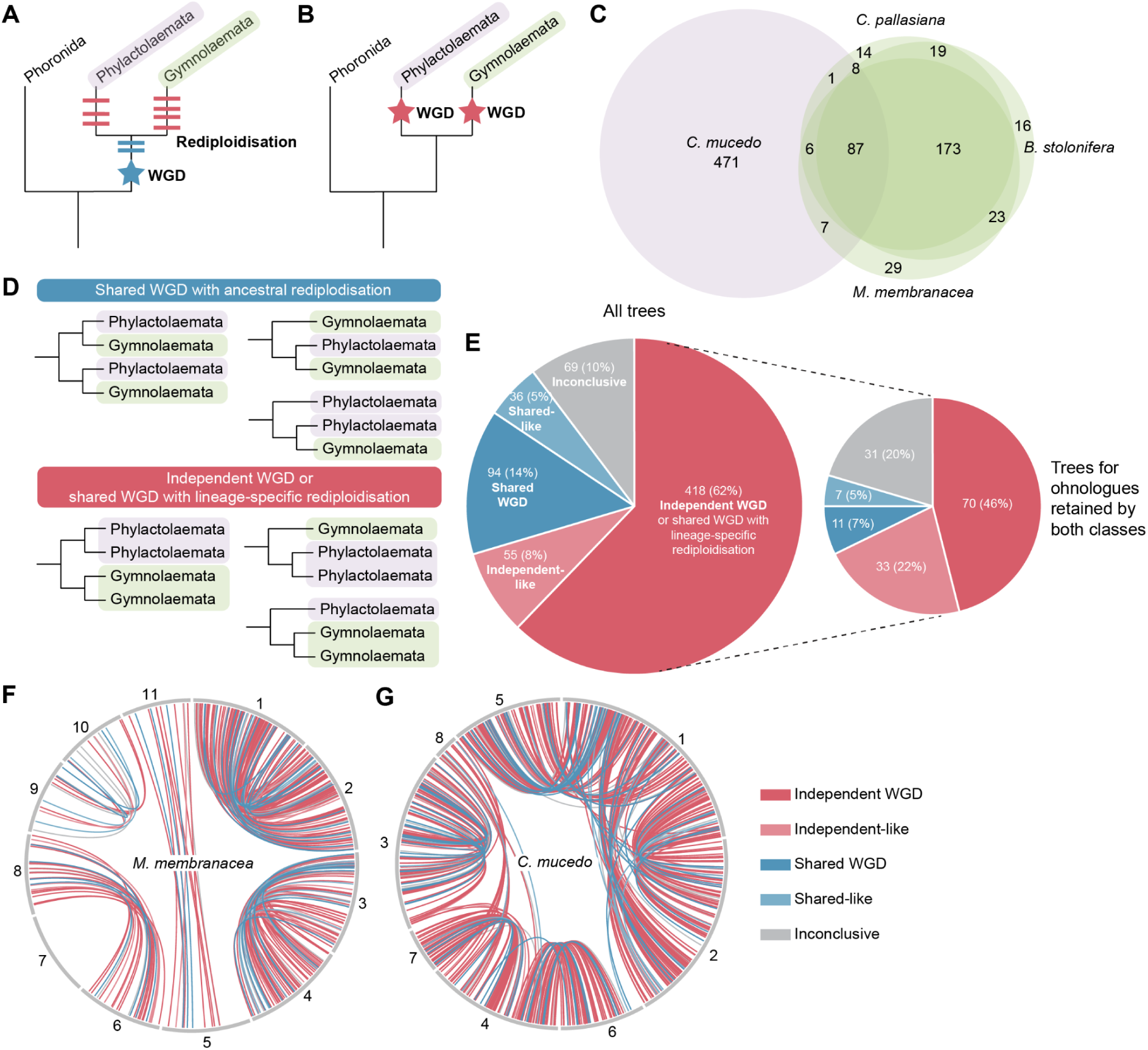
Evidence for a single shared versus multiple independent bryozoan WGDs. A. Hypothesis 1: a single WGD occurred in the ancestor of all bryozoans. Depending on the timing of rediploidisation, duplicates may show a shared WGD topology (blue) or independent WGD topology (red). B. Hypothesis 2: independent WGDs occurred in phylactolaemates and gymnolaemates. All duplicates will show an independent WGD topology. C. Number of ohnologues retained in duplicate in four bryozoans - the phylactolaemate *C. mucedo* (purple) and three gymnolaemates (green). D. Classification of gene tree topologies supporting a shared WGD with ancestral rediploidisation or independent WGDs/shared WGD with lineage-specific rediploidisation. E. Proportion of gene trees with each topology. Large chart represents all gene trees. Smaller chart represents only gene trees containing ohnologues retained in duplicate in both phylactolaemates and gymnolaemates. F. Distribution of ohnologue pairs with each topology around the genome of the gymnolaemate *M. membranacea*. G. Distribution of ohnologue pairs with each topology around the genome of the phylactolaemate *C. mucedo*.

We therefore constructed maximum likelihood gene trees for all ohnologue pairs retained in phylactolaemates, gymnolaemates, or both classes. Tree topologies were classified into those supporting a single shared WGD, multiple independent WGDs, and neither scenario (Fig. 5D, Supplementary Fig. 11, Supplementary Table 19). Independent-like and shared-like topologies were assigned when trees did not perfectly conform to the independent or shared topologies but were closer to one than the other. At this point, we note that rediploidisation—the point at which distinct ohnologous genes form and can start to diverge in sequence—can be asynchronous across the genome. This means that some genes may rediploidise near instantaneously while others may not rediploidise for tens or even hundreds of millions of years post WGD and after speciation events, which complicates interpretation of gene tree topologies^18,139–144^. For instance, the gene tree for an ohnologue pair deriving from a shared WGD, but undergoing lineage-specific rediploidisation (in this case, delayed rediploidisation that post-dates the phylactolaemate-gymnoleaemate split) would be indistinguishable from that expected under independent WGDs^18,139^ (Fig. 5A). Across all ohnologue pair gene trees, 70% recover independent gene duplications in phylactolaemates and gymnolaemates (supporting either independent WGDs or a shared WGD with lineage-specific rediploidisation), 19% show a shared gene duplication event (supporting a shared WGD with shared rediploidisation), while 10% recovered neither topology (Fig. 5E). Consistent with this, a stricter set of gene trees permitting only ohnologues retained in duplicate in both phylactolaemates and gymnolaemates exhibit 68% independent gene duplications and 12% shared gene duplication (Fig. 5E). Phylogenetic analysis of our manually annotated Hox cluster genes indicate that they duplicated/rediploidised independently in each lineage (Supplementary Fig. 10).

Although the shared gene duplication trees are far less frequently observed, this topology is never expected to be recovered if there are independent WGD events. The distribution of gene tree topologies thus supports a single, shared WGD ancestral to all Bryozoa but with most rediploidisation occurring in a lineage-specific manner after the divergence of Phylactolaemata and Gymnolaemata. However, given the unusually small number of ohnologues originating at this proposed WGD time in the bryozoan ancestor, we performed further checks on the validity of these ohnologue pairs. A key indicator of whether these two distinct topologies are real is whether the genes supporting each scenario are located in distinct blocks or regions within the genome^18^. We therefore tracked the location of ohnologue pairs producing trees with each topology in gymnolaemate and phylactolaemate genomes (Fig. 5F, G). The small number of shared WGD trees makes statistical testing the non-randomness of this distribution difficult, so we focussed on the *C. mucedo* genome, which has retained the most ohnologue pairs, to maximise the number of trees for statistical analysis (Fig. 5F, G). We found that chromosome 3 is enriched for genes with a shared WGD topology (Benjamini-Hochberg-corrected Fisher’s exact test *p* < 0.05) (Supplementary Table 20). This suggests that genes forming trees with the two topologies are not randomly distributed around the genome, supporting them as distinct gene sets with different rediploidisation histories following a shared WGD.

Finally, we questioned whether trees recovering either topology were more likely to be the product of systematic errors. To test this, we measured seven metrics for each alignment and/or tree: (1) alignment length, (2) proportion of parsimony-informative sites^145^, (3) relative composition variation^146^, (4) saturation^147^, (5) treeness^146^, (6) average bootstrap support, and (7) standard deviation of branch length heterogeneity^148^. Trees recovering the independent WGD topology (independent and independent-like) were better for three metrics (saturation, average bootstrap support, and standard deviation of branch length heterogeneity), trees recovering a shared WGD topology (shared and shared-like) were better for one metric (proportion of parsimony-informative sites), and there was no difference between the two topologies for three metrics (alignment length, relative composition variation, treeness) (Supplementary Fig. 12, Supplementary Table 21). This does not provide a clear indication that trees supporting either topology are more likely to be the product of systematic errors in tree construction. Overall, we have demonstrated that bryozoans from both Phylactolaemata and Gymnolaemata have a WGD. The most parsimonious explanation for this is a single, common WGD with extensive lineage-specific rediploidisation, but until genome sequencing data from more species becomes available we cannot rule out independent WGDs having occurred in each lineage.

## Discussion

### Ancient WGD in the phylum Bryozoa

Ancient WGDs played a fundamental role in the evolution of plants^4–7,149^ and vertebrates^1,9–14^, generating new alleles and ultimately thousands of new genes that could be recruited to modified roles. In contrast, the importance of WGD to deep invertebrate lineages continues to be uncertain^25^. Our study reveals that many and probably all bryozoans underwent a WGD event in their evolutionary history. In the phylactolaemate *C. mucedo*, an abundance of paired, duplicated, collinear gene blocks distributed across all chromosomes provides incontrovertible evidence of WGD, despite the lack of a temporal peak of gene duplication visible to *K_s_*analysis^95^. In gymnolaemates, cross-phylum synteny comparisons reveal 2:1 and 4:1 ratios of ALGs compared to closely related outgroups such as phoronids, brachiopods, nemerteans, annelids, and molluscs. The chromosomal distribution of gene duplicates within the genomes supports a single WGD as the likely cause of these ratios.

The inference of an ancient WGD over fission is further supported by the monocentric nature of the chromosomes of both phylactolaemates and gymnolaemates^150,151^: fissions of monocentric chromosomes are rare^152^ because they generate an acentric chromosome fragment that requires the de novo evolution of a functional neocentromere to be stably inherited. The fission of every chromosome in the genome exactly once to give a consistent 2:1 chromosome ratio therefore seems especially unlikely. In this light, the absence of clear, duplicated, gene blocks in gymnolaemates is most likely explained by WGD followed by extensive gene loss.

An important question is whether there was a single WGD event inherited by these two bryozoan lineages, or whether independent WGDs occurred in each class. Almost three quarters of gene trees support independent WGDs while around 20% support a single shared event (the rest are ambiguous). At face value, this suggests independent WGDs are more likely—but what explains the highly supported trees with a topology reflecting a shared event? As we noted above, this topology is never expected to be recovered if there are independent WGD events. One explanation is that the WGD was shared but is being partially obscured by lineage-specific rediploidisation. Indeed, there is a precedent from both Acipenseriformes fishes and snow carp for independent WGD events in sister lineages being erroneously inferred due to asynchronous rediploidisation^140,153–158^ when in reality it was a single, shared WGD^18^. Gene duplicates only begin to diverge following rediploidisation, and rediploidisation can occur asynchronously in different parts of a genome^139^. Therefore, if some genes rediploidise before, and some after, a speciation event, genes from different genomic regions will have differing evolutionary histories and tree topologies^139^. The observed mix of tree topologies is consistent with a single WGD shared by gymnolaemates and phylactolaemates with a small amount of the genome rediploidising prior to the split of these lineages, and most of the genome rediploidising independently in each group.

If there was a single ancient WGD shared by all bryozoans, this predicts that not only *C. mucedo* but all phylactolaemates should display signatures of WGD. Though *C. mucedo* remains the only phylactolaemate genome available, Saadi et al.^121^ performed a survey of 16 phylactolaemate species’ transcriptomes (representing over 18% of species of this small clade). In their dataset, the Hox cluster duplication is found in 15/16 species, with the missing one likely due to assembly quality. Taking Hox cluster duplication as a marker of WGD, that provides strong evidence that the WGD that we observe in *C. mucedo* is indeed phylactolaemate-wide. Overall, when taking into account parsimony and gene tree topologies, we deem a single, bryozoan-wide WGD the most likely scenario but cannot with current data fully rule out the possibility of independent WGDs in phylactolaemates and gymnolaemates.

If phylactolaemate and gymnolaemate bryozoans indeed share a paleopolyploidy event, it would make Bryozoa the first animal phylum known to have an ancestral, founding WGD. This would add bryozoans to seed plants and vertebrates as speciose, ‘evolutionarily successful’ lineages underpinned by a WGD. Given the deep divergence of phylactolaemates and gymnolaemates, it would be amongst the oldest paleopolyploidies described. The recent discovery of unequivocal stenolaemate (generally accepted as the sister group to gymnolaemates) bryozoan fossils from the early Cambrian places a conservative absolute minimum bound on any bryozoan-wide WGD at 514 MYA^70^. This would place it earlier than recent estimates of the 2R event ancestral to all jawed vertebrates, while positioning it contemporaneous with or older than vertebrate 1R^9,10^. Future estimates of WGD age based on the divergence of high-confidence, early-rediploidising, ohnologues should help establish a more specific age interval for the bryozoan WGD.

Functional redundancy means that many duplicates rapidly return to single-copy post-WGD^159^, while the rate and extent to which this occurs likely varies across WGD events^160^. In bryozoans, we estimate only 5–10% of genes to have been retained in duplicate post-WGD. In comparison, the retention rate of duplicates post WGD is around 15%^161^ in teleosts (WGD 235–350 MYA)^162–165^, 7.5–20.5% in arachnopulmonates (450 MYA)^33^, and 39–58% in *Paramecium* (320 MYA)^160^. The retention of so few WGD-derived duplicates in bryozoans raises the question of whether paleopolyploidy had any lasting impact on their long-term evolution. For example, it is tantalising that the occurrence of WGD likely coincides with the major transition of bryozoans to coloniality, which presumably required a substantial increase in the complexity of gene regulation^166^.

We contend that even WGDs with very low rates of gene retention could be evolutionarily important by breaking up gene clusters and removing regulatory constraints^167^. We found bryozoan genes retained in duplicate to be enriched in transcription factors, similar to results from vertebrates^13,168^, angiosperms^169–171^ and fungi^172^. Transcription factors are dosage-sensitive because duplications and losses can disrupt stoichiometric ratios and cause dysregulation of gene expression^173,174^. They are therefore expected to be retained post-WGD at an elevated rate but are rarely duplicated as part of small-scale duplications^160,175,176^. This gives significant potential for the rewiring of gene regulatory networks post-WGD, potentially facilitating functional innovation and the evolution of lineage-specific novelties.

In bryozoans, *rfx*, a key transcriptional regulator of ciliogenesis^125–127^, has been retained in duplicate following WGD, and further genes involved in the regulation, construction, and functioning of cilia were also retained at an elevated rate. Using RNA-seq data, we found that orthologues of ciliary genes duplicated by WGD are preferentially expressed in the lophophore in adult bryozoans, implying that duplicates created at the WGD were recruited to roles in lophophore cilia. Why might this have been favoured in bryozoans, when phoronids and brachiopods also possess ciliated lophophores? One possibility is that it contributes to bryozoans’ ability to produce multiciliate cells^177^, which may represent an adaptation to the reduction of lophophore size in bryozoans^178^ in comparison to the larger, monociliate lophophores of phoronids and brachiopods. Bryozoan cilia are also highly diverse: in adults, water-pumping lateral cilia on the lophophore beat in a coordinated rhythm to generate feeding currents ^137^. Food particles are then collected by stiff laterofrontal cilia in a process known as ciliary sieving^138^, and finally transported towards the mouth by a third group of cilia, the frontal cilia^179^ (though these are lacking in stenolaemates^180^). Bryozoan cyphonautes larvae are also ciliated planktotrophic filter-feeders, using a mechanism that is highly distinct from that of adults^181^.

### Rampant gene loss and the detection of cryptic WGDs

It is logical that most methods used for WGD detection^52^ centre on the presence or organisation of duplicated genes. Synonymous substitution (*K_s_*) distributions test for peaks in duplicates of a specific age^47–49^, intra-genomic synteny analysis searches for duplicated collinear blocks^99,100,182,183^, and gene tree-species tree reconciliation identifies bursts of gene family duplications on a specific branch^7,50–52^. However, our analyses add to a growing weight of evidence that extensive gene loss and/or long time periods since a WGD limit the applicability of such methods to paleopolyploidy events^8,50^. Taking *K_s_* distributions as an example, *K_s_* saturation can create artificial peaks at older *K_s_* values, real WGD peaks are flattened by stochastic variation as they get older, and duplicates arising from small-scale duplications can obscure WGD-derived peaks when there are few true WGD duplicates, as is frequently the case following paleopolyploidy^48,50,184^. The lower limit of *K_s_* distributions to detect a WGD has been estimated at 10% of duplicate gene retention^49^ and distinguishing ancient WGDs from background duplications is problematic and risks becoming subjective^171^. Moreover, recent work increasingly points to rediploidisation—which is the true point at which a gene can be considered duplicated—occurring asynchronously across a genome over the course of tens of millions of years^18,139–144^. This makes approaches that assume WGD creates a burst of duplicates of an identical age fundamentally unreliable for WGD inference. The present bryozoan case exemplifies this point in particular, since most ohnologues diverging after speciation means that targeting genes duplicated on the proposed WGD branch would yield in the region of an order of magnitude fewer ohnologues with which to detect an already cryptic WGD.

Macrosynteny is considered to be the most robust method to study ancient WGDs^8,25,53,56,115,185,186^, for instance being used to disentangle the number and timing of WGDs in vertebrates/cyclostomes^9,10,55^ and flowering plants^8^ and demonstrate the absence of WGD in lepidopterans^115^. Cross-species orthologous synteny has particularly strong potential because, in the absence of significant interchromosomal rearrangements, it could in principle detect many-to-one chromosomal ratios even if all genes have returned to single copy. We propose that integrating ALGs may add further power to such analyses by accommodating interchromosomal rearrangements, since many-to-one ALG ratios should be preserved even in the face of extensive chromosome fusions (though not fissions). This may increase the distances at which such comparisons can be made, and is critical for the applicability of this framework in invertebrates, where chromosome fusions are pervasive^58,187^. Bryozoans provide a particularly compelling case study of this approach, demonstrating how ALG-based synteny can extend the detection of ancient WGDs to highly rearranged invertebrate genomes. In gymnolaemates, we uncovered a cryptic WGD with ohnologue retention estimated at around 5% that was invisible to *K_s_* distributions^95^ and paralogous synteny. Only a combination of orthologous synteny with bilaterian ALG information and intraspecies macrosyntenic gene distributions was able to reveal the presence of a WGD. These results raise the possibility that cryptic WGDs are being systematically overlooked, and therefore that the contribution of ancient WGDs to lineages like invertebrates is currently underappreciated. With genome sequencing efforts now increasingly turning to lesser-studied phyla^75,188–193^ and ever more sophisticated methods for WGD detection, we expect that further cryptic WGDs will soon emerge from within the invertebrates.

## Methods

### Dataset curation and genome annotation

We assembled a dataset of all available chromosome-level bryozoan genomes (*n* = 9)^89–93,96,194^, and supplemented this with genomes from the two phyla most closely related to bryozoans, Phoronida and Brachiopoda (Supplementary Table 1). Only chromosome-level genomes were considered for analysis because synteny-based WGD detection requires highly contiguous assemblies. Published annotations^75,95,96,191^ were used for all except two of these genomes. The two remaining genomes, those of *Parasmittina trispinosa* and *Schizoporella japonica*, were downloaded in soft-masked format from NCBI Datasets^195^ and annotated using BRAKER3 (v3.0.8)^103,196–204^. For *S. japonica*, transcriptomic data (SRA dataset ERR12245645) were combined with the metazoan protein set from OrthoDB (v11)^205^ as hints for annotation. For *P. trispinosa*, OrthoDB (v11)^205^ protein hints alone were used in the absence of RNA data. Annotation performance was assessed using BUSCO (v6.0.0) with the Lophotrochozoa_odb12 dataset^112,206^. To standardise gene prediction quality and filter annotations for spurious proteins and repetitive sequences, we performed self-blasts of each species’ proteome with DIAMOND (v2.1.16)^207^ and removed protein sequences which had more than 50 hits at --evalue 1e-10.

### Phylogenomic analysis

Single-copy orthologues were identified with Orthofinder (v3.1.0)^208^. Each orthogroup was then aligned with MAFFT (v7.525)^209^ in mode L-INS-i and trimmed with trimAl (v1.5.rev0)^210^ with mode “-automated1”, optimised for maximum likelihood phylogenetics. Trimmed alignments were then concatenated with PhyKIT (v2.1.2)^211^. The tree was constructed in IQ-TREE (v3.0.1)^110^ with 1000 UFBoot2 ultrafast bootstraps^212^ and the Q.PFAM+F+R7 model as selected by ModelFinder^111^.

### Intra-species synteny analysis

Two complementary approaches were used to identify blocks of conserved synteny within genomes (paralogons). First, duplicated collinear gene blocks were identified with MCScanX (v1.0.0)^107^ using a block size of 2 and maximum gaps threshold of 50, then plotted as Circos diagrams^213^ with shinyCircos (v2.0)^214^. Second, the collinearity search algorithm of i-ADHoRe (v3.0.01)^108^ was run in wgd (v2.0.38)^215^ with ‘wgd syn’ after the identification of homologues using DIAMOND^207^ with ‘wgd dmd’.

### Inter-species synteny analysis

SyntenyFinder (v1.0)^95^ was used to characterise macrosyntenic relationships across genomes. SyntenyFinder first implements Orthofinder (v3.1.0)^208^ to identify orthologues. This Orthofinder-based orthology is then used to determine to which of the 24 bilaterian ancestral linkage groups^26,57^ each gene belongs. SyntenyFinder then combines orthology information, ALG assignments, gene locations from gtf files, and karyotype information from genome fasta files to produce dot plots and ribbon plots, which are constructed using RIdeogram (v0.2.2)^216^.

### Tests for enrichment of gene duplicates on chromosome pairs

First, reciprocal best hits (RBH) were identified in each genome using DIAMOND blastp (v2.1.16)^207^ with the following parameters: --outfmt 6 --max-target-seqs 5 --evalue 1e-70 --no-self-hits --header --ultra-sensitive. A strict e-value was used to restrict the dataset to only very high confidence RBH (average of 952 RBH pairs per species). RBH with both pairs on the same chromosome were eliminated as likely tandem duplicates. We then constructed a contingency table of RBH counts between each chromosome pair and tested for deviations from a random distribution of hits using a chi-squared test. *p*-values were calculated using the pchisq function in the R stats package (v4.3.0)^217^ and multiple tests were accounted for with a Bonferroni correction. Where the global test indicated a significant departure from random pairing, standardised Pearson residuals were used to identify chromosome pairs contributing most strongly to this signal. Standardised Pearson residuals were plotted as heatmaps.

### Analysis of retained ohnologues

To quantify the proportion of ohnologue pairs retained in extant bryozoan genomes, we first used Orthofinder (v3.1.0)^208^ to find orthologues between bryozoans and representatives of their two nearest outgroups, the phoronid *Phoronis australis* and the brachiopod *Lingula anatina*. Putative retained ohnologues were then identified in a two step process. First, orthogroups that were single-copy in phoronids and brachiopods but duplicated (2–5 copies) in a given bryozoan were extracted. Second, these orthogroups were filtered to retain only those with gene pairs on each of two chromosomes identified as putatively ohnologous based on synteny. The proportion of ohnologues retained post-WGD was then calculated as this gene set divided by the total number of orthogroups that were single-copy in phoronids and brachiopods and present in a given bryozoan. This method prevents the erroneous counting of tandem-duplicates as ohnologues, but means that our inferred retention rate may be an underestimate if ohnologues have been translocated between chromosomes.

Overlaps of retained ohnologue sets were then analysed using DeepVenn (beta version)^218^. Gene Ontology (GO) enrichment analysis on retained ohnologue sets was performed using topGO (v2.52.0)^219^ with the ‘classic’ algorithm and Fisher test. GO-mapping and functional annotation of bryozoan protein sets was performed by the orthology-based method of eggNOG-mapper (v2.1.13)^220^ with eggNOG DB (v5.0.2)^221^ and DIAMOND (v2.0.15)^207^. BUSCO duplication counts of bryozoans were compared to species with and without a WGD using BUSCO (v6.0.0) with the Lophotrochozoa_odb12 dataset^112,206^.

### Construction of gene trees

Gene trees were constructed by alignment of protein sequences with MAFFT (v7.525)^209,222^, trimming with trimAl with option -gappyout (v1.5.rev0)^210^, and initial maximum likelihood tree-building using IQ-TREE (v3.1.2)^110^ with 1000 ultrafast bootstraps^212^ and ModelFinder^111^ selection of the optimal model of sequence evolution. This initial tree was then used as a guide tree (option -ft) to construct a final tree with the posterior mean site frequency (PMSF) ^223^ approximation of the CAT site-specific model ^224^ to account for site-specific evolutionary rate heterogeneity within a maximum likelihood framework. Gene trees were rendered in iTOL (v6)^225^. For each tree, the potential for systematic errors driving the topology was measured using metrics calculated with PHYKIT^211^: (1) alignment length, (2) proportion of parsimony-informative sites^145^, (3) relative composition variation^146^, (4) saturation^147^, (5) treeness^146^, (6) average bootstrap support, and (7) standard deviation of branch length heterogeneity^148^. Higher values are preferable for alignment length, proportion of parsimony-informative sites, saturation, treeness and bootstrap support. Lower values are preferable for relative composition variation and standard deviation of branch length heterogeneity.

### Analysis of RNA sequencing data

Ciliary genes were defined as those included in the CiliaCarta database^124^. A one-sided Fisher’s Exact Test was used to test for an enrichment of ciliary genes in the retained ohnologue set of the phylactolaemate *C. mucedo* and gymnolaemates (represented by *M. membranacea*). Our dataset of gene expression in lophophore versus non-lophophore (gut) tissue in the bryozoan *Flustrellidra hispida*^75^ was used to test if bryozoan ciliary genes with retained ohnologues are preferentially expressed in their ciliated feeding organ (lophophore). Salmon (v1.10.0)^226^–derived TMM values were plotted using pheatmap (v1.0.13)^227^. Benjamini-Hochberg-corrected *t*-tests were used to test for a difference between lophophore and non-lophophore expression. *p* < 0.1 was considered significant.

## Data Availability

Gene models for the two bryozoan species annotated in this study have been deposited on Figshare (https://figshare.com/articles/dataset/Bryozoan_genome_annotations/33368995). Gene Trees are provided as part of Supplementary Table 19. The SyntenyFinder pipeline is available on GitHub (https://github.com/symgenoevolab/SyntenyFinder).

## Acknowledgements

The authors were supported by a Biotechnology and Biological Sciences Research Council (BBSRC) Strategic Longer and Larger Award (BB/Z51746X/1) that funds the WGDip Consortium Project (https://www.rediploidisation.org/). A.K.R. was supported by a Royal Society Research Ireland University Research Fellowship (URF\R1\241884) and a University College Dublin Ad Astra Fellowship. We are grateful to Alex de Mendoza, Dan J. Macqueen, Ferdinand Marlétaz, Peter O. Mulhair, Stephan Q. Schneider and members of the WGDip Consortium for insightful discussions. We would also like to thank all attendees of the International Bryozoology Association’s 19^th^ Larwood Symposium, held at the Natural History Museum in Oslo, for their advice, discussions, and encouragement (https://www.bryozoology.org/). We thank Jo Wood of the Tree of Life Programme at the Wellcome Sanger Institute for providing the Hi-C contact map for *S. japonica*, and Patrick Adkins, Freja Azzopardi, John Bishop, Helen Jenkins, Rebekka Uhl and Chris Wood for initial collection and identification of bryozoan specimens for the Tree of Life Programme. This research utilised Queen Mary University of London’s Apocrita HPC facility, supported by QMUL Research-IT (http://doi.org/10.5281/zenodo.438045).

