## Supplementary Material for "Cryptic whole-genome duplication in bryozoans"

#### This PDF file includes

Supplementary Discussion 1–2

Supplementary Figures 1–12

Legends for Supplementary Tables 1–21

References

#### **Supplementary Discussion 1: Lineage-specific rearrangements in *Schizoporella japonica***

In our dataset of nine bryozoans, eight have chromosome numbers between 8 and 12. *S. japonica* is an outlier, with 19 chromosomes, suggesting additional lineage-specific rearrangements. We first questioned whether this is genuine or the result of assembly error, but a Hi-C contact map supports the division of the *S. japonica* genome into 19 chromosomes (Supplementary Fig. S4).

We then turned to riparian plots to understand how the *S. japonica* karyotype emerged from the ancestral gymnolaemate karyotype of 11 chromosomes. This shows that *S. japonica* has undergone 10 fissions and two fusions to give its 19 chromosomes (Supplementary Fig. S5). The possibility of an additional whole-genome duplication (WGD) in *S. japonica* can be ruled out because riparian plots comparing its genome to its nearest relative show that each *S. japonica* chromosome corresponds to only one chromosome arm in *W. subatra*, not a full chromosome (Supplementary Fig. S5).

Furthermore, just prior to submission of this manuscript, another *Schizoporella* genome, *S. errata*, was released on NCBI (GCA\_980783175.1). *S. errata* possesses the standard gymnolaemate karyotype of 11 chromosomes, suggesting that the rearrangements that have occurred in *S. japonica* are very recent and are not even shared by the full genus.

**Supplementary Discussion 2: Ancient fissions and whole-genome duplications leave different signatures in the distribution of gene duplicates**

In this work, we use the distribution of gene duplicates across chromosomes within a genome to distinguish between two hypotheses: WGD versus chromosome fission. This is based on the assumption that only WGD and not fission will produce an excess of duplicates across chromosome pairs (Main Text Fig. 3A, B). To validate this assumption, we compared the distribution of duplicates following known fission events to the putative WGDs.

Firstly, the evolutionary history of gymnolaemate chromosomes includes an ancient fission event<sup>1</sup> shared by all currently sequenced gymnolaemates (Supplementary Fig. S7A). Following sequential fusions that combined genes from 18 of the 24 bilaterian ancestral linkage groups (ALGs)<sup>2</sup> onto one chromosome, this chromosome then fissioned to produce two daughters. Our data suggests that, at the WGD, each of these daughters was then duplicated, leaving four chromosomes carrying these 18 ALGs. This explains the presence of a 4:1 ratio of these chromosomes compared to outgroups, while most others show a 2:1 ratio. Within this quartet of chromosomes, each is related to the other three by either fission or WGD, but never both. We therefore compared the enrichment of genes across chromosomes related by fission versus duplication. In each case, chromosomes related by WGD contain a strong enrichment of gene duplicates while those produced by fissions do not (Supplementary Fig. S7B). This validates our contention that WGD but not fission will produce chromosome pairs with an excess of gene duplicates.

Second, as described in Supplementary Discussion 1, the genome of *S. japonica* contains nine lineage-specific fission events that are not found even in close relatives. This offers us a near-perfect test case, with many examples of chromosomes derived from fission or WGD in the same genome (Supplementary Fig. S8A). Pearson standardised residuals for chi-square tests for enrichment of gene duplicates between pairs of chromosomes that were derived from WGD are significantly higher than those produced by fissions ($p = 1.68 \times 10^{-5}$) (Supplementary Fig. S8B–D). This again supports the assumption that WGD but not fissions result in pairs of chromosomes sharing an excess of duplicate genes.

Overall, these two examples show that neither ancient fission events occurring before the WGD nor more recent lineage-specific fission events result in an enrichment of duplicates across daughter chromosome pairs. This confirms that the enrichments observed are not the product of fissions, and therefore must be derived from WGD.

### Supplementary Figures 1–12

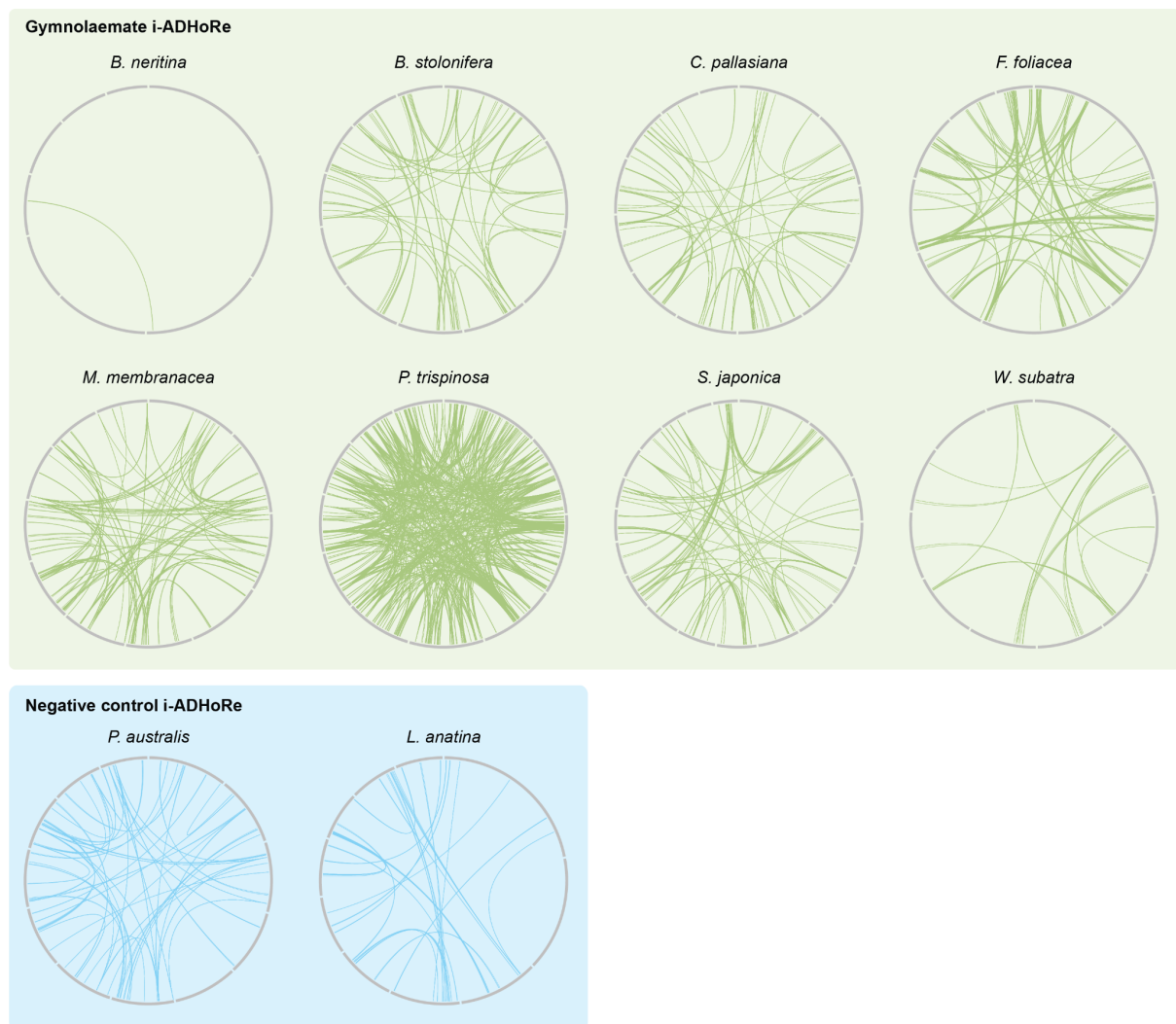

**Supplementary Fig. 1. Distribution of i-ADHoRe paralogs in gymnolaemate genomes**

Circos plots showing the distribution of duplicated collinear gene blocks (paralogons) in the genomes of eight gymnolaemate bryozoans, the phoronid *Phoronis australis*, and the brachiopod *L. anatina*, as determined by i-ADHoRe<sup>3</sup>. Grey bars are chromosomes. Green or blue lines connect the position of duplicated collinear gene blocks (paralogons). Only cross-chromosome paralogs are shown.

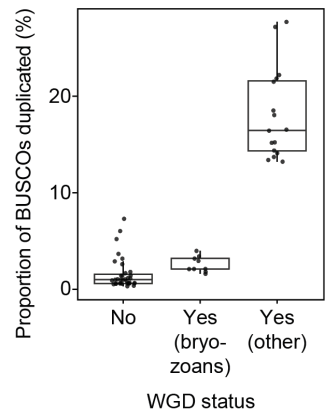

**Supplementary Fig. 2. Proportion of duplicated BUSCOs in spiralian with and without a WGD.**

Plot shows proportion of duplicated Benchmarking Universal Single-Copy Orthologs (BUSCOs)<sup>4</sup> for genomes with no WGD, certain WGD, and the nine chromosome-level bryozoan genomes. Bryozoan genomes are closer in BUSCO duplication rate to genomes with no WGD.

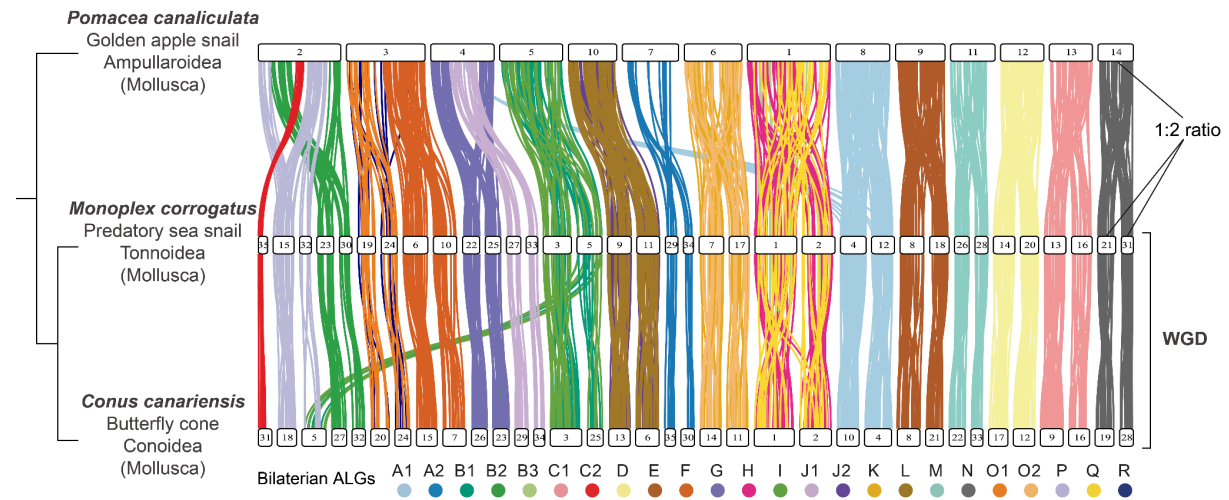

**Supplementary Fig. 3. Riparian plot showing the 2:1 chromosome ratio for a described WGD in caenogastropod molluscs.**

The caenogastropod molluscs *Monoplex corrogatus* and *Conus canariensis* share a WGD that is not found in outgroups like *Pomacea canaliculata* (golden apple snail)<sup>5,6</sup>. As a result, there is a 1:2 ratio when chromosomes are compared using the 24 bilateral ancestral linkage groups (ALGs)<sup>2</sup>. White bars represent chromosomes. Vertical ribbons connect the position of orthologous genes, coloured by their bilateral ALG.

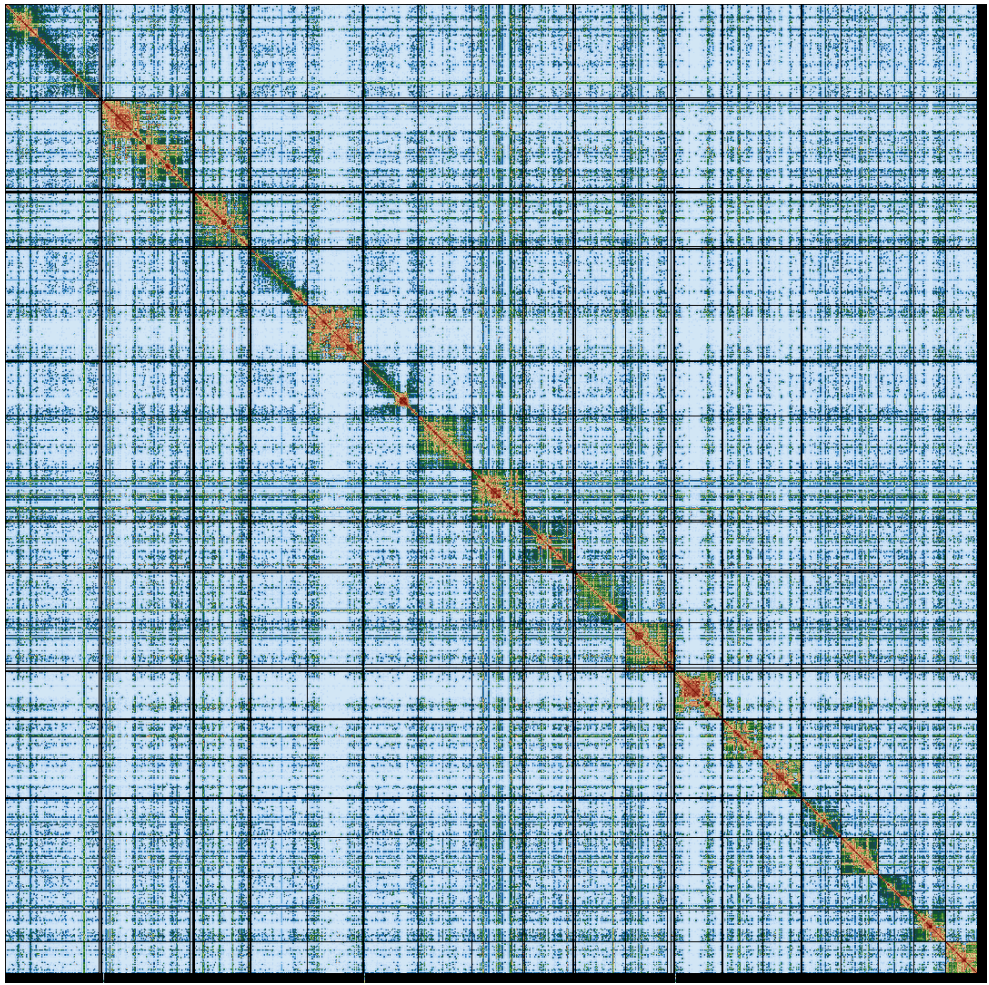

**Supplementary Fig. 4. Hi-C contact map for the bryozoan *Schizoporella japonica*.**

The contact map shows that the chromosome fissions observed in the *S. japonica* genome are genuine and not the product of assembly errors. The map is shown for the primary haplotype tzSchJapo1.1 (GCA\_965278275.1). Figure provided by the Darwin Tree of Life Project at the Wellcome Sanger Institute.

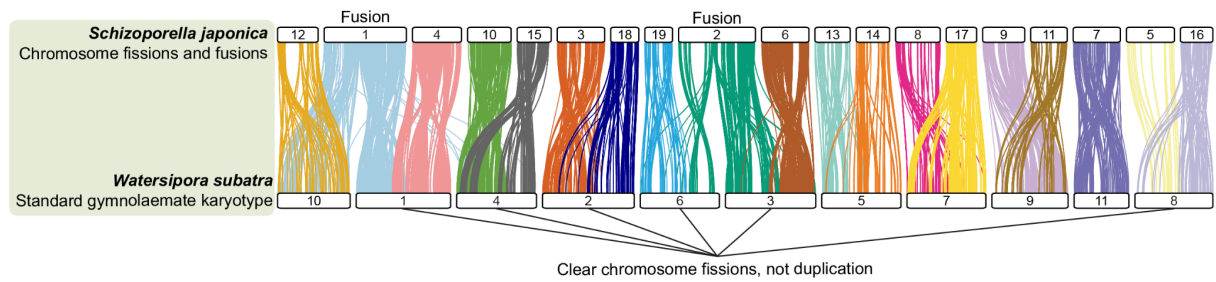

**Supplementary Fig. 5. Riparian plot for *S. japonica* and *W. subatra*.**

White bars represent chromosomes. Vertical ribbons connect the position of orthologous genes, coloured by their chromosome in the *S. japonica* genome. The plot demonstrates that the additional chromosomes in *S. japonica* are the product of fissions, not an additional round of WGD. This is because each *S. japonica* chromosome contains genes from only half the corresponding chromosome in the nearest outgroup, *W. subatra*.

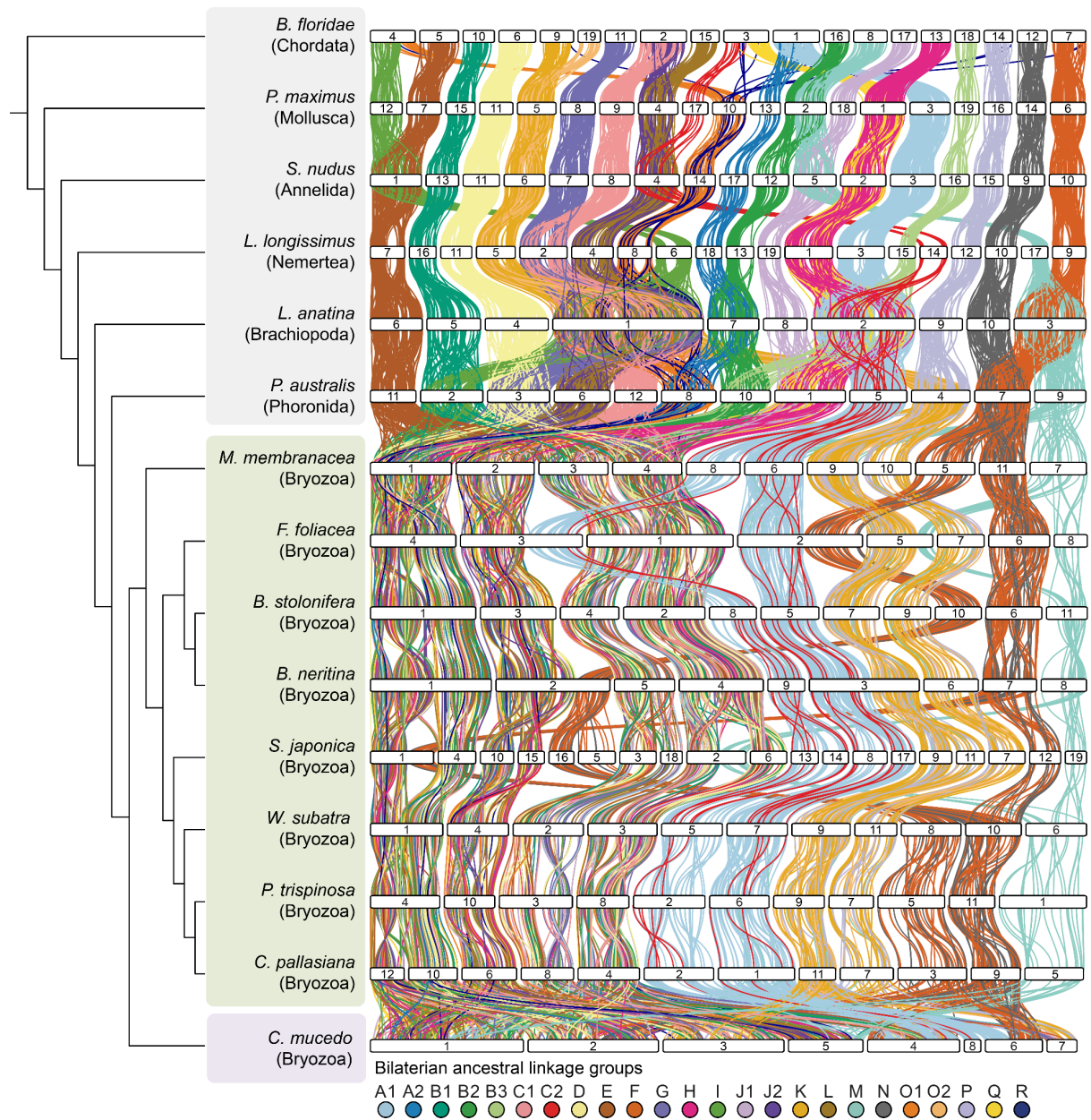

**Supplementary Fig. 6. Riparian plot for nine bryozoan genomes and representatives of six other animal phyla.**

White bars represent chromosomes. Vertical ribbons connect the position of orthologous genes, coloured by their bilaterian ALG<sup>2</sup>. In outgroup phyla, each ALG is found on one chromosome. In most gymnolaemate bryozoans, each ALG is found on two (A1, C2, F, K, N, O2, P) or four (A2, B1, B2, B3, C1, D, E, G, H, I, J1, J2, L, O1, Q, R) chromosomes. This is suggestive of either one or two rounds of WGD or one or two fissions of each ancestral chromosome. In *S. japonica*, ALGs are distributed across three (K, O2, P) four (A1, C2, F, N) or eight (A2, B1, B2, B3, C1, D, E, G, H, I, J1, J2, L, O1, Q, R) chromosomes due to additional fissions and fusions (see Supplementary Discussion 1). ALG M is distributed across many chromosomes in all gymnolaemates, suggesting it was dispersed prior to WGDs/fissions.

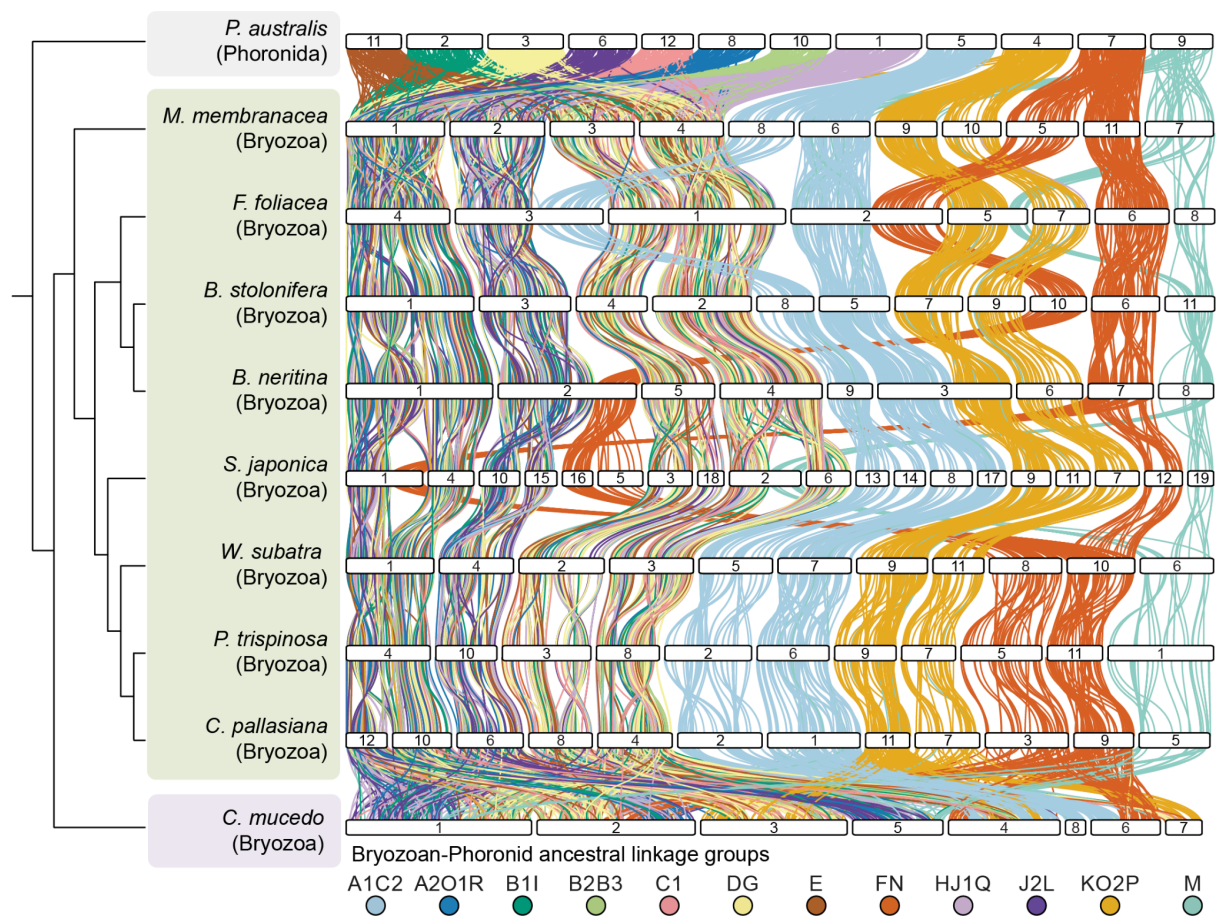

**Supplementary Fig. 7. Riparian plot for nine bryozoan genomes and *P. australis*, coloured by bryozoan-phoronid ALGs.**

White bars represent chromosomes. Vertical ribbons connect the position of orthologous genes, coloured by their bryozoan-phoronid ALG<sup>7</sup>. In *P. australis*, each ALG is found on one chromosome. In most gymnolaemate bryozoans, each ALG is found on two (A1C2, FN, KO2P) or four (A2O1R, B1I, B2B3, C1, DG, E, HJ1Q, J2L) chromosomes. This is suggestive of either one or two rounds of WGD or one or two fissions of each ancestral chromosome.

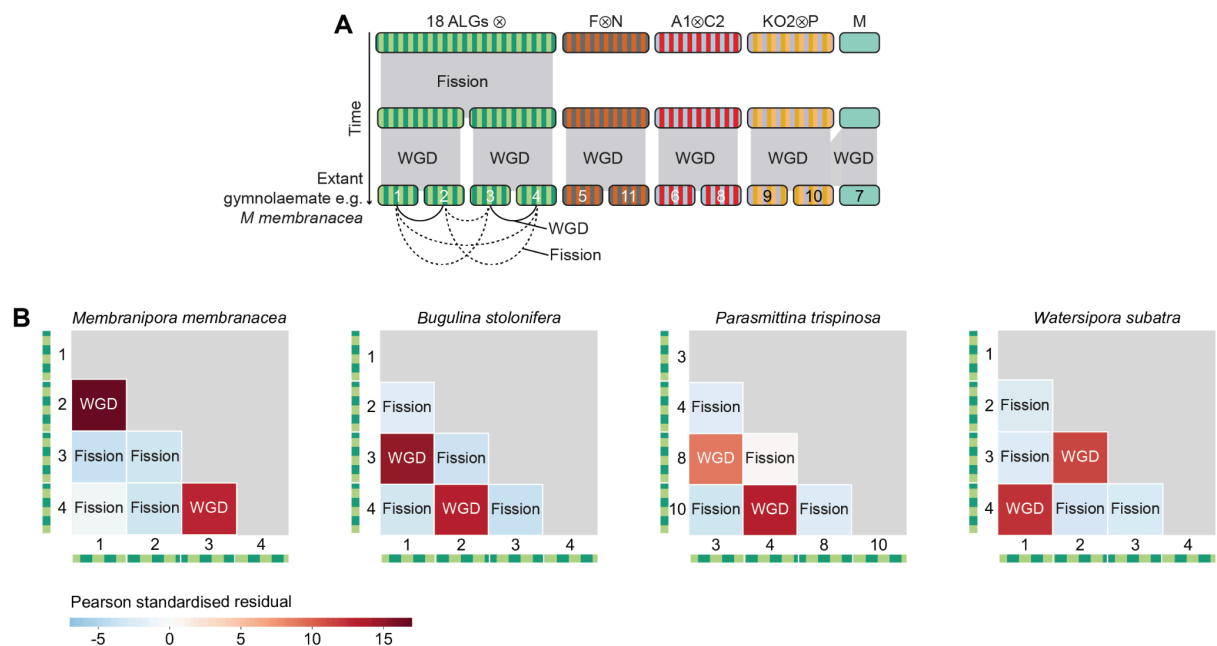

**Supplementary Fig. 8. Fissions and WGD leave different distributions of gene duplicates in gymnolaemate bryozoans.**

A. Reconstruction of karyotype evolution in gymnolaemate bryozoans. The chromosome that is the product of many fusions and contains 18 of the 24 bilaterian ALGs first fissioned into two. Each of these two chromosomes was then duplicated by the WGD, leaving four daughter chromosomes each carrying genes from all of the 18 ALGs. Each of these four chromosomes is related to the other three either by fission or by WGD.

B. Chromosomes related by WGD show a strong enrichment of gene duplicates but those related by fission do not. Heatmaps show chromosome pairs enriched (red) and depleted (blue) in duplicates. Only species in which these chromosomes have not been involved in subsequent rearrangements are shown.

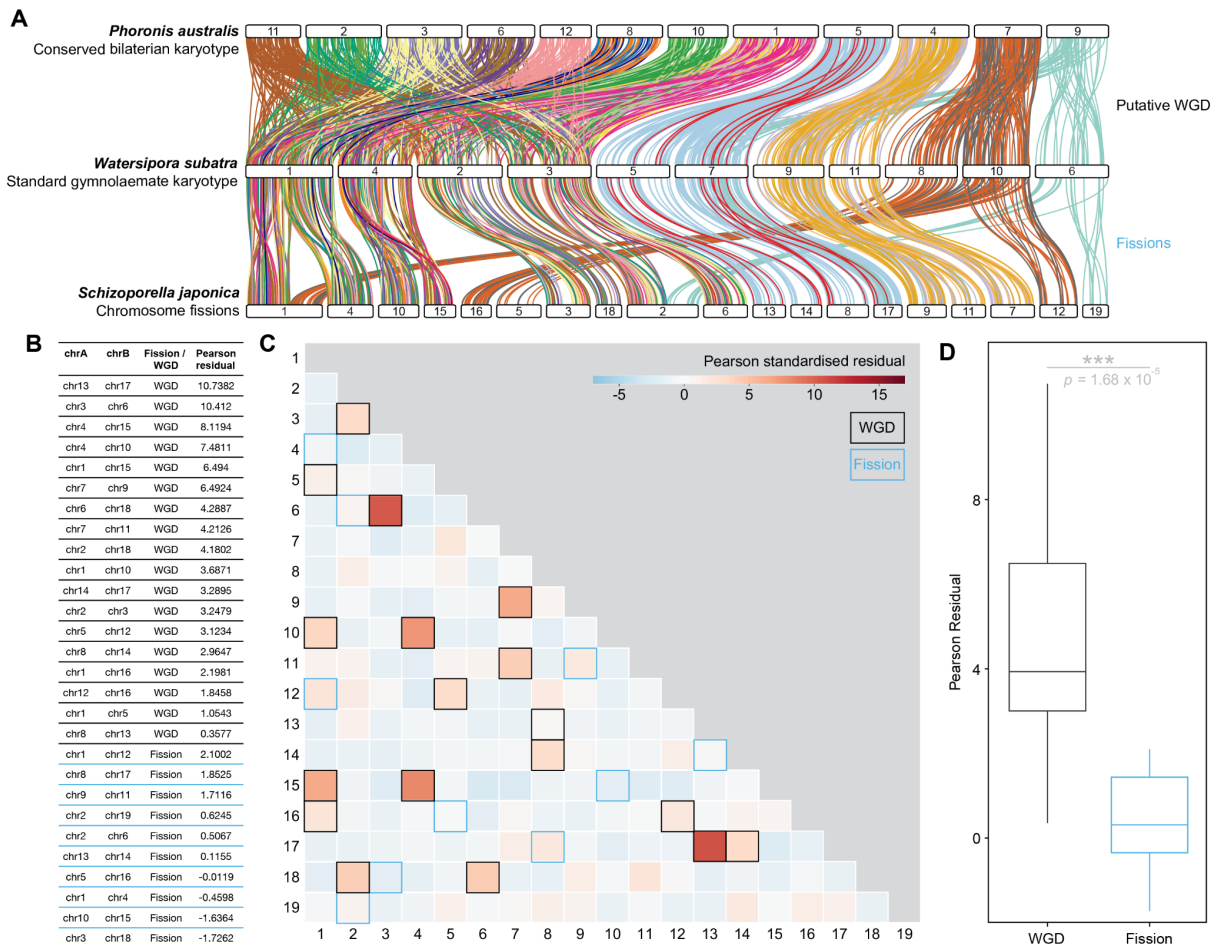

**Supplementary Fig. 9. Fissions and WGD leave different distributions of gene duplicates in *S. japonica*.**

A. Riparian plot for *S. japonica*, *W. subatra* and *P. australis*. White bars represent chromosomes. Vertical ribbons connect the position of orthologous genes, coloured by their bilaterian ALG.

B. Pearson residual for enrichment of gene duplicates on pairs of chromosomes each derived from a WGD or chromosome fission event.

C. Heatmap showing chromosome pairs enriched (red) and depleted (blue) in duplicate gene pairs. Each chromosome pair is represented by one square. Black and blue boxes mark chromosome pairs derived from WGD or fission, respectively.

D. Boxplot showing Pearson residuals for chromosome pairs derived from WGD or chromosome fission events. Residuals from WGD are significantly higher than chromosome fission (Welch's *t*-test, $p = 1.68 \times 10^{-5}$).

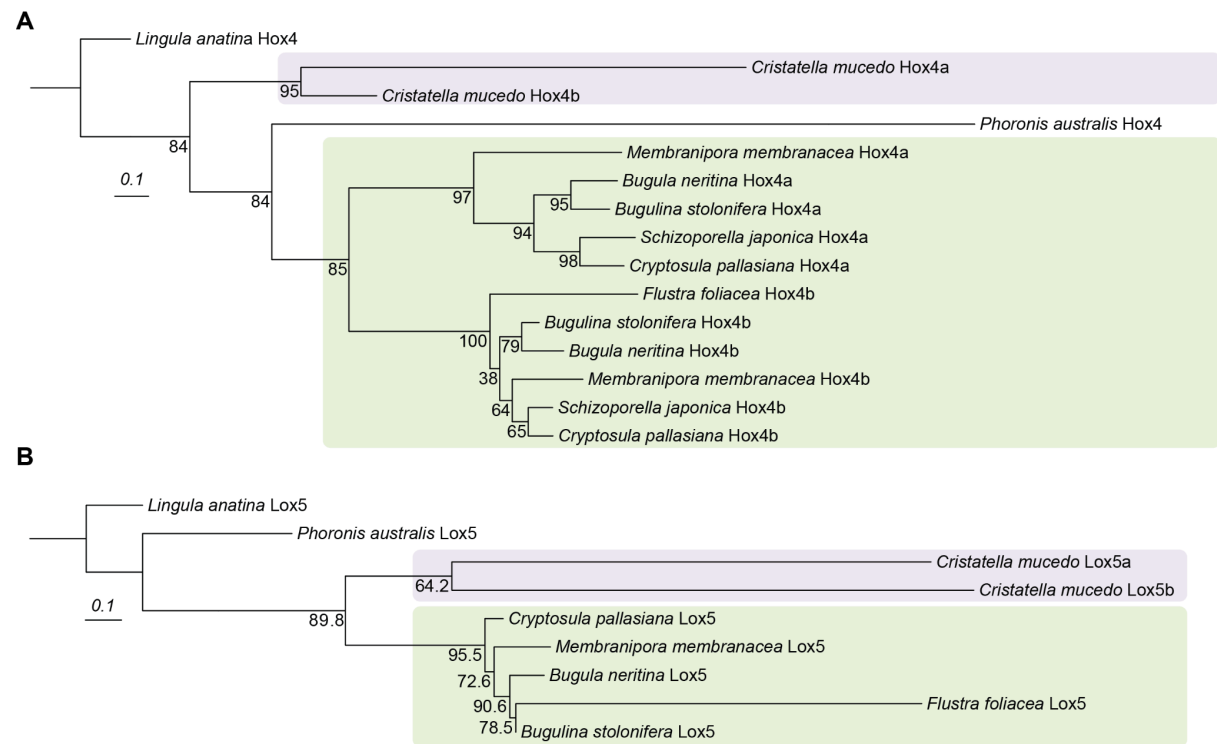

**Supplementary Fig. 10. Gene trees of bryozoan duplicated Hox proteins Hox4 (A) and Lox5 (B).**

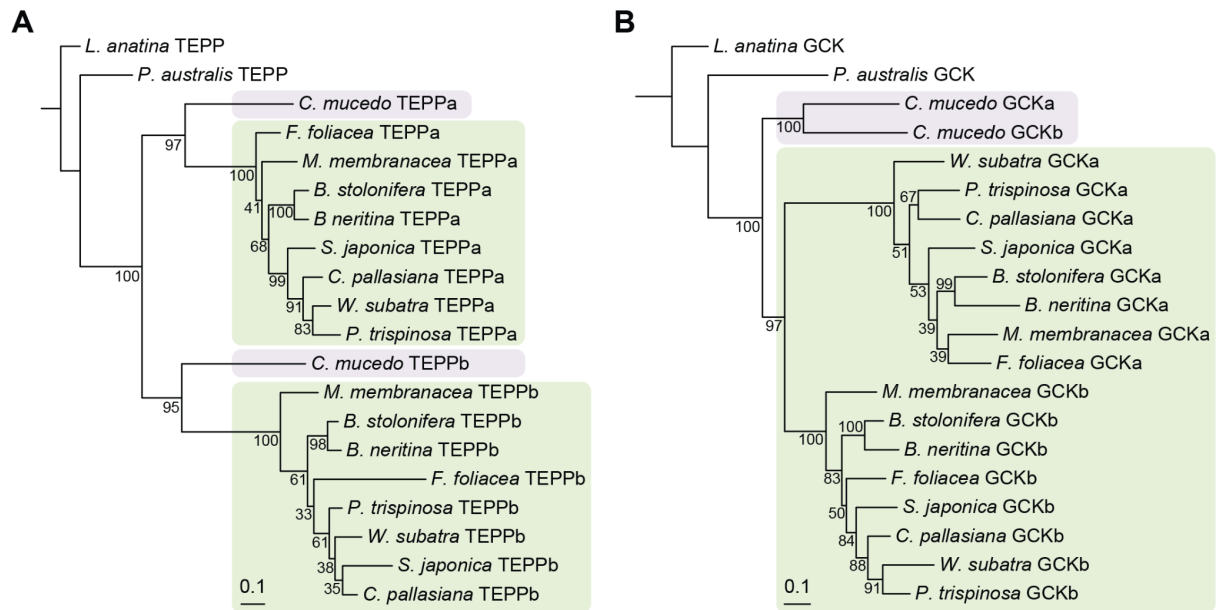

**Supplementary Fig. 11. Examples of gene trees supporting shared (A) versus independent duplication (B).**

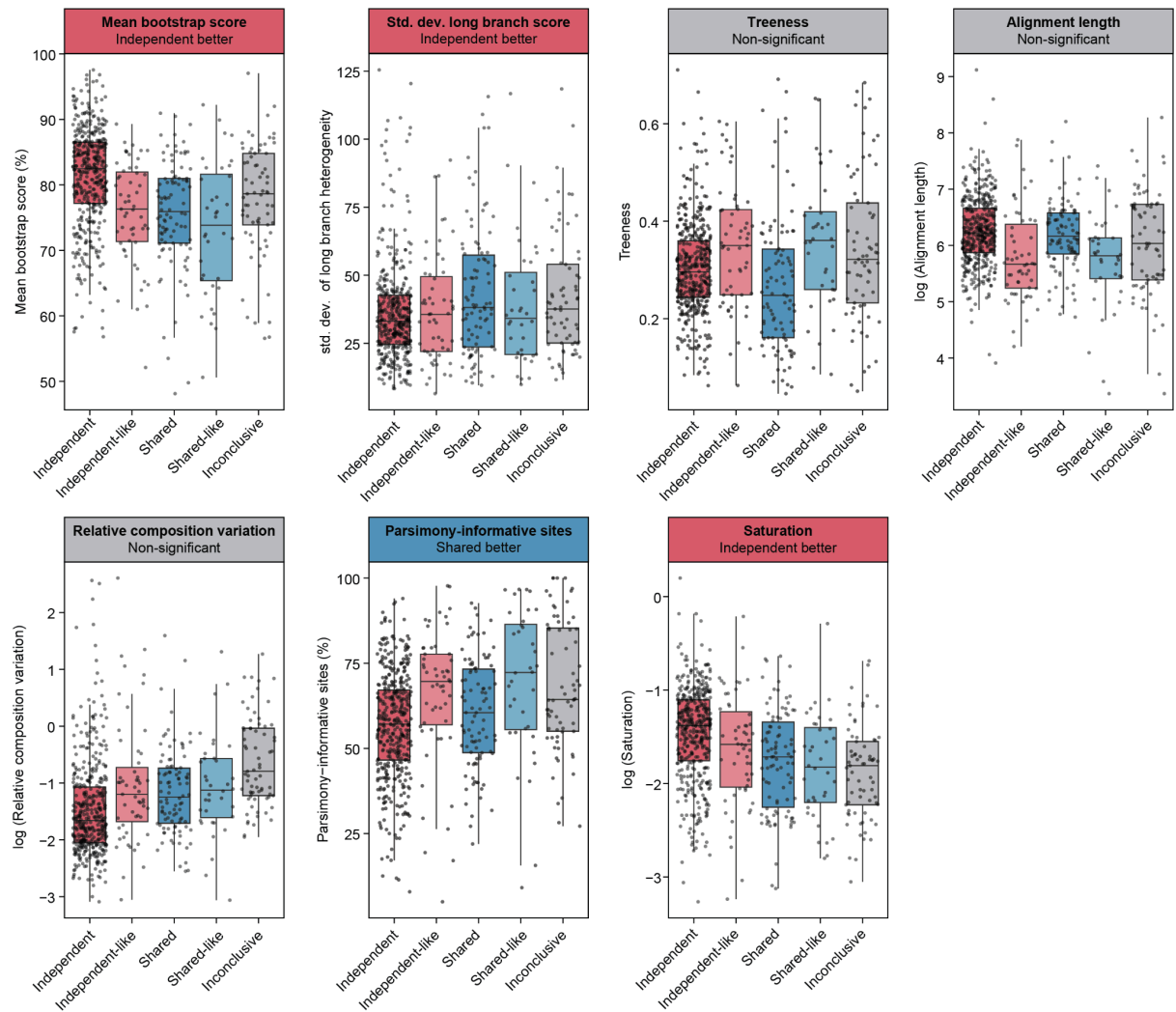

**Supplementary Fig. 12. Likelihood of systematic errors in alignments/trees supporting different topologies.** Data for Welch's two-sample *t*-tests available as Supplementary Table 21.

**Legends for Supplementary Tables 1–21**

**Supplementary Table 1.** Genomic dataset used in this work including three-letter species codes.

**Supplementary Table 2.** Duplication rate of BUSCO genes for various spiralian with and without WGD.

**Supplementary Table 3.** Pearson residuals for Chi-squared tests for chromosome fissions versus duplications.

**Supplementary Table 4.** Chi-squared tests for enrichment of gene duplicates between chromosome pairs.

**Supplementary Table 5.** Ohnologue retention rate following WGD in each bryozoan genome.

**Supplementary Table 6.** Retained ohnologues in *C. mucedo*.

**Supplementary Table 7.** Eggnog-mapper annotation of *C. mucedo* genes.

**Supplementary Table 8.** Gene ontology molecular function for *C. mucedo* retained ohnologues.

**Supplementary Table 9.** Gene ontology biological process for *C. mucedo* retained ohnologues.

**Supplementary Table 10.** Retained ohnologues in *M. membranacea*.

**Supplementary Table 11.** Eggnog-mapper annotation of *M. membranacea* genes.

**Supplementary Table 12.** Gene ontology molecular function for *M. membranacea* retained ohnologues.

**Supplementary Table 13.** Gene ontology biological process for *M. membranacea* retained ohnologues.

**Supplementary Table 14.** Ciliary genes duplicated in bryozoans.

**Supplementary Table 15.** Fisher's exact test results for enrichment of ciliary genes in retained ohnologue sets.

**Supplementary Table 16.** Trimmed Mean of M-values (TMM)-normalised expression matrix for *Flustrellidra hispida* gut and lophophore samples.

**Supplementary Table 17.** Eggnog-mapper annotation of *F. hispida* genes.

**Supplementary Table 18.** Expression of ciliary genes in *F. hispida* RNA-seq data.

**Supplementary Table 19.** Tree topologies and Newick trees for each duplicated ohnologue.

**Supplementary Table 20.** Distribution of trees with each topology in the *C. mucedo* genome.

**Supplementary Table 21.** Welch's two-sample *t*-tests for differences in mean bootstrap score, standard deviation of branch length heterogeneity, treeness, alignment length, relative composition variation, parsimony-informative sites, and saturation for trees with independent(-like) and shared(-like) topologies.
